# Single-cell transcriptomics reveals targeted modulation of inflammatory repertoire by SOCE blockers

**DOI:** 10.1101/2025.04.28.651042

**Authors:** Andreas Stephanou, Madhav Mantri, Divya Shankaranarayanan, Carol Li, Mila Lagman, Jenny Xiang, Chendong Pan, Yanjie Sun, Thangamani Muthukumar, Khaled Machaca, Iwijn De Vlaminck, Manikkam Suthanthiran

## Abstract

Store-operated calcium entry (SOCE) plays a critical role in regulating intracellular calcium signaling and is essential for immune cell functions. SOCE blockade with pyrazole derivative BTP2 has been explored as an anti-inflammatory strategy in preclinical models and Zegocractin (CM4620) is being investigated in Phase 2 clinical trials as an immunoregulatory agent. However, the mode of action and differential effects of SOCE blockade on diverse immune cell types remain largely unknown, limiting the precision of current therapeutic applications. Here, we used multiplexed single-cell RNA sequencing to investigate the effects of two prototypic SOCE blockers, BTP2 and CM4620, on polyclonally-stimulated, normal human peripheral blood mononuclear cells (PBMCs). The data revealed that SOCE blockade suppresses the expression of cytotoxicity-associated genes in CD8□ effector T cells and natural killer (NK) cells, restoring them to levels comparable to those in unstimulated cells. Strikingly, SOCE blockade preserved activation-induced expression of anti-inflammatory genes in CD4□ regulatory T cells, maintaining their tolerance phenotype even after SOCE blockade. These findings suggest that SOCE blockers modulate immune responses with greater selectivity than conventional immunosuppressants by reducing cytotoxicity while preserving tolerance-associated pathways, highlighting their potential for managing immune-mediated conditions, including organ transplantation.

**KEY POINTS:**

- SOCE blockers BTP2 and CM4620 reduce cytotoxic gene expression in CD8□ T cells and NK cells.
- SOCE blockade preserves anti-inflammatory programs in CD4□ Tregs, supporting selective immune modulation.

## INTRODUCTION

Calcium (Ca^2^□) is an important second messenger and is critical for cellular function^1^. The activation of membrane phospholipase C, followed by the generation of inositol triphosphate (IP□) and its binding to receptors on the endoplasmic reticulum (ER), triggers the release of Ca^2^□ from the ER into the cytosol^2^. Depletion of Ca^2^□ from the ER induces Stromal Interaction Molecule (STIM) proteins to undergo conformational changes and cluster at ER–plasma membrane (PM) junctions, where they recruit Orai channel family members to mediate store-operated calcium entry (SOCE), resulting in a sustained increase in cytosolic Ca^2^□ ^3,4^.

Calcium channel blockers suppress immune functions by disrupting calcium homeostasis, thereby reducing the risk of transplant rejection or managing autoimmune conditions where excessive immune responses can be harmful^5,6^. SOCE blockade results in an altered activation of transcription factors such as CREB and NF-κB, which regulate genes involved in various cellular processes critical to immune function such as activation, proliferation, differentiation, cytokine production and apoptosis^4,7^. Calcium signaling regulates the activation of transcription factors such as nuclear factor activated T cells (NFAT) which is crucial for T cell activation and cytokine release^8,9^. Calcium signaling has also been found to be essential in macrophages for the regulation of phagocytosis as well as modulation of the production of pro-inflammatory cytokines/chemokines following microbial stimuli^10^. Given the multifaceted role of calcium in various immunological functions, it is important to precisely understand which aspects of the immune system are impacted by SOCE blocking and to elucidate the underlying molecular mechanisms.

Zegocractin or CM4620, a SOCE blocker that is undergoing phase 2 clinical trial as an anti-inflammatory agent, selectively inhibits the calcium influx mediated by ORAI1, the pore-forming subunit of the calcium release-activated calcium (CRAC) channel^11,12^. CM4620 has shown promise in clinical trials for its ability to selectively target immune cells, thereby providing a novel approach to immune suppression with potentially fewer side effects compared to traditional immunosuppressive drugs^13^. Another small molecule, YM-58483 or BTP2, has been reported to inhibit multiple components within the CRAC channel pathway, namely, STIM1 which is the ER calcium sensor, and ORAI1 which is the pore-forming subunit of the CRAC channel. Additionally, BTP2 has also been shown to affect TRPC channels, which are part of the transient receptor potential channel family involved in calcium entry. Despite targeting multiple components of SOCE, BTP2 remains a potent SOCE inhibitor and has demonstrated efficacy in preclinical studies for its ability to suppress immune responses; however, BTP2 has not progressed to clinical trials for human use^14–17^. Despite increasing insights into the wide-ranging effects of SOCE inhibitors such as BTP2 and CM4620, their specific molecular impact on various immune cell types remains unclear.

Advances in high-throughput single-cell technologies have enabled the study of different cellular processes and pathways in diverse cell types^18,19^. In this study, we utilized multiplexed single-cell RNA sequencing to investigate and compare the effects of BTP2 and CM4620 on PBMCs from healthy donors stimulated with phytohemagglutinin (PHA). Our analysis revealed that SOCE blockade using either BTP2 or CM4620 suppressed the expression of cytotoxicity-associated genes in CD8□ effector T cells and natural killer (NK) cells, restoring them to levels comparable to those in unstimulated PBMCs. Importantly, we find that these treatments spared the activation-induced expression of anti-inflammatory genes in CD4□ regulatory T cells, which maintained their tolerance phenotype even after SOCE blockade. These findings highlight the role of SOCE blockers in modulating inflammation while maintaining immune tolerance.

## METHODS

### Healthy volunteers

Peripheral venous whole blood was obtained from healthy volunteers after obtaining informed written consent. The research project #1208012870 was approved by the Weill Cornell Medicine Institutional Review Board. The authors confirm that all research was performed in accordance with relevant guidelines/regulations, and the research involving human research was performed in accordance with the Declaration of Helsinki.

### PBMC Isolation and Stimulation

Peripheral blood mononuclear cells (PBMCs) were isolated from healthy volunteers using Ficoll-Paque™ density gradient centrifugation. PBMCs (1×10□ cells/mL) were incubated under various conditions, including stimulation with PHA and treatment with BTP2 or CM4620 for 16 hours. Control samples were treated with DMSO. Details of the procedure are provided in the **Supplemental Methods**.

### RNA Isolation and Quantitative PCR

Total RNA was extracted from PBMCs using the RNeasy Mini Kit (Qiagen), followed by quantification and quality assessment. Reverse transcription and quantitative PCR (RT-qPCR) were performed to measure absolute mRNA copy numbers. Primer sequences and additional details are included in the

### Supplemental Methods. Bulk and scRNA Sequencing

For bulk RNA sequencing, RNA samples were treated with DNase before library preparation using the TruSeq RNA Library Prep Kit (Illumina) and sequenced on a NovaSeq 6000 system. Single-cell RNA sequencing (scRNA-seq) was performed using the 10X Genomics 3’ Gene Expression platform, with library preparation and sequencing conducted at the Weill Cornell Medicine Genomics Core. Additional processing details are provided in the **Supplemental Methods**.

### Bulk and scRNA-seq Data Processing and Analysis

Bulk RNA-seq reads were aligned to the human genome (GRCh38) using STAR, and gene expression counts were processed with DESeq2 for normalization and differential expression analysis^20–24^. scRNA-seq data were preprocessed using Scanpy, with quality control, normalization, clustering, and annotation based on canonical immune cell markers^25^. Details of filtering criteria, clustering parameters, and computational tools are included in the **Supplemental Methods**.

### Gene Expression Similarity, Effect Size Estimation, and Enrichment Analysis

To assess similarity between treatment conditions, Pearson correlation, Euclidean distances, and cosine similarity metrics were computed. To quantify the effect of treatments on gene expression profiles, Cohen’s d was calculated for key gene modules, including cytotoxicity and tolerance scores. Gene set enrichment analysis (GSEA) was performed using GSEApy to identify pathways associated with cytotoxicity and tolerance^26–29^. Further details are provided in the **Supplemental Methods**.

## RESULTS

### RNA sequencing of PHA-stimulated human PBMCs treated with SOCE blockers

To investigate the effect of SOCE channel blockers on diverse immune cell types including at the single cell level, we treated PHA-stimulated primary peripheral blood mononuclear cells (PBMCs) from healthy human volunteers with two SOCE channel blockers, BTP2 or CM4620 (**Methods, Fig. 1A**). As controls, we used unstimulated and untreated cells, PHA-stimulated but untreated cells, unstimulated but BTP2 treated cells and unstimulated but CM4620-treated cells from the same donors. To investigate the transcriptional effects of SOCE channel blockers and the cell-type specificity of these effects on diverse immune cell types, we performed both bulk and single-cell RNA sequencing on human PBMCs across six conditions: (i) DMSO (Control), (ii) BTP2, (iii) CM4620, (iv) PHA, (v) PHA + BTP2, (vi) PHA + CM4620. (**Fig. 1A)**. Supplementary Figure 1 (**Supp. Fig. 1**) shows quality control for bulk-RNA-seq samples across different donors and treatment conditions. We analyzed the bulk RNA-seq data to understand the aggregated impact of SOCE blockers on bulk transcriptomes of PHA-stimulated PBMCs. Dimensional reduction and visualization of bulk transcriptomes using Principal Component Analysis (PCA) for samples across all six treatment conditions revealed three main groups. Control PBMCs, PBMCs treated with BTP2 alone, and PBMCs treated with CM4620 alone formed a single cluster, PBMCs stimulated with PHA formed a separate cluster, and the third cluster consisted of PBMCs stimulated with PHA and treated with BTP2 and PBMCs stimulated with PHA and treated with CM4620 (**Fig. 1B**). We performed hierarchical clustering on batch corrected RNA-seq data and found that the samples clustered based on treatment conditions and not based on individual volunteers, confirming that the source of PBMCs had a minimal impact (**Fig. 1C**). Supp. Table 1 provides the normalized base mean counts for all genes across the six experimental conditions.

**Figure 1:**
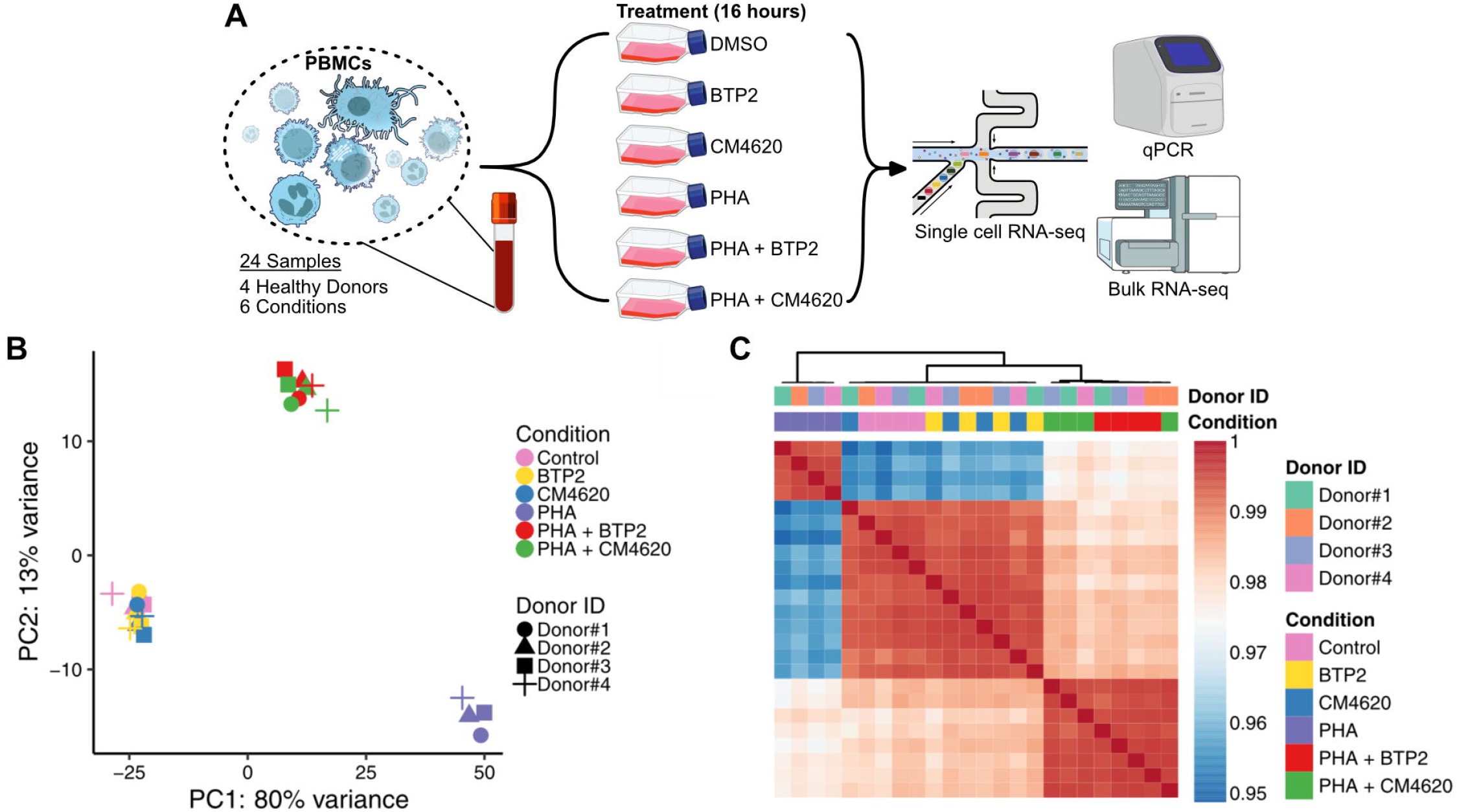
RNA sequencing of PHA-stimulated human PBMCs treated with SOCE blockers. **A)** Experimental workflow for RNA-seq of PHA-stimulated human PBMCs treated with SOCE blockers BTP2 and CM4620. **B)** Principal component analysis (PCA) plot of PBMC transcriptomes from four healthy donors colored by treatment conditions. **C)** Heatmap showing hierarchical clustering of PBMC transcriptomes from four healthy donors colored by treatment conditions.

We performed differential gene expression analysis (DGEA) on PBMCs stimulated with PHA and compared them to control PBMCs. DGEA revealed a significant upregulation of 3519 genes in PHA-stimulated cells (log_2_ fold-change > 1.0, adjusted p-value < 0.01, **Supp. Fig. 2A)**. Among these, 94 genes showing the highest fold-changes (log_2_ fold-change > 5.0, adjusted p-value < 0.01) in PHA-stimulated cells were associated with ontology terms such as cellular response to interleukin-1, response to chemokines, neutrophil chemotaxis, leukocyte proliferation and differentiation, and T cell differentiation (**Supp. Fig. 2B**). Next, to understand the effect of SOCE blockers on PHA-stimulated cells, we compared significantly altered genes from PBMCs + PHA vs. PBMCs + PHA + BTP2, and significantly altered genes from PBMCs + PHA vs. PBMCs + PHA + CM4620. DGEA revealed a downregulation of 1364 and 1339 genes in PHA-activated cells when treated with BTP2 and CM4620, respectively (log_2_ fold-change < -1.0, adjusted p-value < 0.01, **Supp. Table 1**). Comparison of DGEA results for PHA-stimulated cells treated BTP2 or CM4620 revealed a high correlation between changes in transcriptional changes induced by the two SOCE entry blockers BTP2 and CM4620 (R = 0.95, p-value = 2.2e-16, **Supp. Fig. 2C**). This was further supported by almost no differences observed in a direct comparison between PHA-stimulated cells treated with BTP2 and PHA-stimulated cells treated with CM4620 (**Supp. Fig. 2D, Supp. Table 1**). Our global gene expression analysis identified that SOCE blockade with both BTP2 and CM4620 mostly spares activation-induced expression of anti-inflammatory genes (e.g., less than 50% inhibition of activation induced expression of genes for TGFB1, IL10, and CTLA4) whereas the induced expression of proinflammatory genes such as IFNG (97% inhibition by BTP2 and 97% inhibition by CM4620) and cytopathic genes such as GZMB (83% inhibition by BTP2 and 84% inhibition by CM4620 were markedly inhibited (**Supp. Fig. 3, Supp. Table 1**).

We utilized an orthogonal platform, the RT-qPCR assay, to validate the differential impact of BTP2 and CM4620 on PHA-induced alterations in gene expression. Our measurement of absolute mRNA copy number using the RT-qPCR assay confirmed RNA-seq identified differential impact on gene expression. As observed with RNA-seq, the expression level of mRNAs for the T cell receptor CD3E and T cell subtypes CD4 and CD8A were not significantly altered by either BTP2 or CM4620 (**Supp. Fig. 4**). As found using RNA seq, ITGAE mRNA was significantly downregulated by PHA activation, and this down regulation was significantly reversed by both BTP2 and CM4620 (**Supp. Fig. 4**). In accord with RNA Seq findings, both BTP-2 and CM4620 significantly and markedly suppressed PHA-induced expression of mRNAs encoding T cell costimulatory molecules CD27 and CD28, as well as mRNAs for IL-2, CD25, proinflammatory IFNG, IFNG induced chemokines CXCL9 and CXCL10, and cytopathic proteins perforin and granzyme B. Also, as found by RNA seq, neither BTP2 nor CM4620 altered the PHA-induced increase in the expression of mRNA for prototypic immunosuppressive cytokine TGFB1. There was a significant but modest suppression of mRNAs for IL10 and CTLA4 (**Supp. Fig. 4**). Collectively, these results demonstrate that both BTP2 and CM4620 markedly block pro-inflammatory gene expression while relatively sparing the expression of genes encoding anti-inflammatory proteins.

### scRNA sequencing of PHA-stimulated human PBMCs treated with SOCE blockers

To investigate the cell-type specificity of the effects of SOCE channel blockers on diverse immune cell types, we performed single-cell RNA sequencing on human PBMCs across all six treatment conditions (**Fig. 1A, Supp. Fig. 5A-B)**. After removing cells with low mRNA counts and potential cell doublets, we detected a mean of 3,471 to 5,222 transcripts across donors (**Supp. Fig. 5A**). Analyzing the number of genes detected and the number of transcripts across conditions revealed a higher number of distinct genes, as well as a greater total number of transcripts, in PBMCs activated with PHA (**Supp. Fig. 5B**). Interestingly, PHA-activated PBMCs treated with BTP2 or CM4620 exhibited lower transcript diversity and transcript counts compared to PHA-activated cells but still displayed higher values than control PBMCs (**Supp. Fig. 5B**). Pearson correlation analysis across samples within each donor revealed high correlations, typically ranging from 0.92 to 1.0, between technical replicates indicating strong reproducibility within each condition (**Supp. Fig. 6**). Clustering patterns revealed that replicates from the same treatment condition group tightly together, further emphasizing consistency (**Supp. Fig. 6**).

We analyzed single-cell transcriptomes from PBMCs (Control) condition alone to characterize the cellular heterogeneity in unstimulated and untreated PBMCs (**Supp. Fig. 7A-C**). We detected distinct transcriptomic clusters for various immune cell types such as B cells, CD14^+^ classical monocytes, CD16+ non-classical monocytes, dendritic cells, CD8^+^ T cells, CD4^+^ T cells, NK cells and plasma cells (**Supp. Fig. 7A-C**). For the CD8^+^ T cell subtypes, we detected distinct clusters for naive and effector cells, with naive CD8^+^ cells transcriptionally more similar to CD4^+^ T cells than to the effector CD8^+^ T cells (**Supp. Fig. 7A**).

To analyze the effect of SOCE blockers on unstimulated PBMCs, we analyzed unstimulated PBMCs treated with BTP2 or CM4620 with control PBMCs. Unsupervised clustering on single-cell transcriptomes from these three conditions revealed distinct clusters for all immune cell types containing cells derived from all three conditions (**Supp Fig. 7D-F**). Differential gene expression analysis on BTP2 or CM4620 treated PBMCs compared to control PBMCs for each cell type showed an upregulation on genes associated with ontology terms such as regulation of viral processes, regulation of innate immune responses, and regulation of defense response for both BTP2 and CM4620 (**Supp. Fig. 8A-D**). When treated with BTP2 or CM4620, an upregulation of genes associated with innate immune response was observed for many cell types with NK cells, monocytes, and CD4 T cells showing the highest increase in the expression of interferon-stimulated genes (ISGs) such as IFI6, ISG15, MX1, XAF1, and IFI44L compared to control PBMCs (**Supp. Fig. 8A-D, Supp. Table 2-3**).

We analyzed single-cell transcriptomes across all six conditions together to understand the effect of BTP2 and CM4620 on PHA-stimulated cells. We clustered the transcriptomes from these six conditions to label populations of all immune cell types. We observed separation of cells from treatment conditions within clusters representing all major cell types including CD8^+^ Naïve T cells, CD8^+^ Effector T cells, B cells, CD14 monocytes and CD16 monocytes (**Fig. 2A-C**). Interestingly, CD8^+^ Naïve T cells, CD4^+^ T cells, as well as CD4^+^ T-reg cells separated into three distinct clusters based on treatment conditions: Cluster 1 containing unstimulated and untreated cells, and unstimulated and BTP2 or CM4620-treated cells, Cluster 2 containing PHA-stimulated cells, and Cluster 3 containing PHA-stimulated and BTP2 or CM4620-treated cells (**Fig. 2A & 2B**). To quantitatively assess the effect of PHA activation and the effect of BTP2 and CM4620 on PHA-stimulated cells, we performed differential gene expression analysis on individual cell types to compare PHA-stimulated cells to control unstimulated cells (**Supp. Fig. 9A & Supp. Table 4**), and to compare stimulated cells treated with SOCE blockers to stimulated but untreated PBMCs (**Supp. Fig. 9B-C & Supp. Table 5-6**). To assess the effect of PHA activation and the effect of BTP2 and CM4620 on PHA-stimulated cells across the entire transcriptome, we next estimated pairwise Pearson correlations for all cell types across all six conditions. Correlation analysis revealed groups of cell types for almost all cell types except CD4^+^ T cells, CD8^+^ Naïve T cells, and CD4^+^ T-reg cells, suggesting that these cell types showed the biggest changes in transcription on PHA activation and after drug treatment. (**Fig. 2D**).

**Figure 2:**
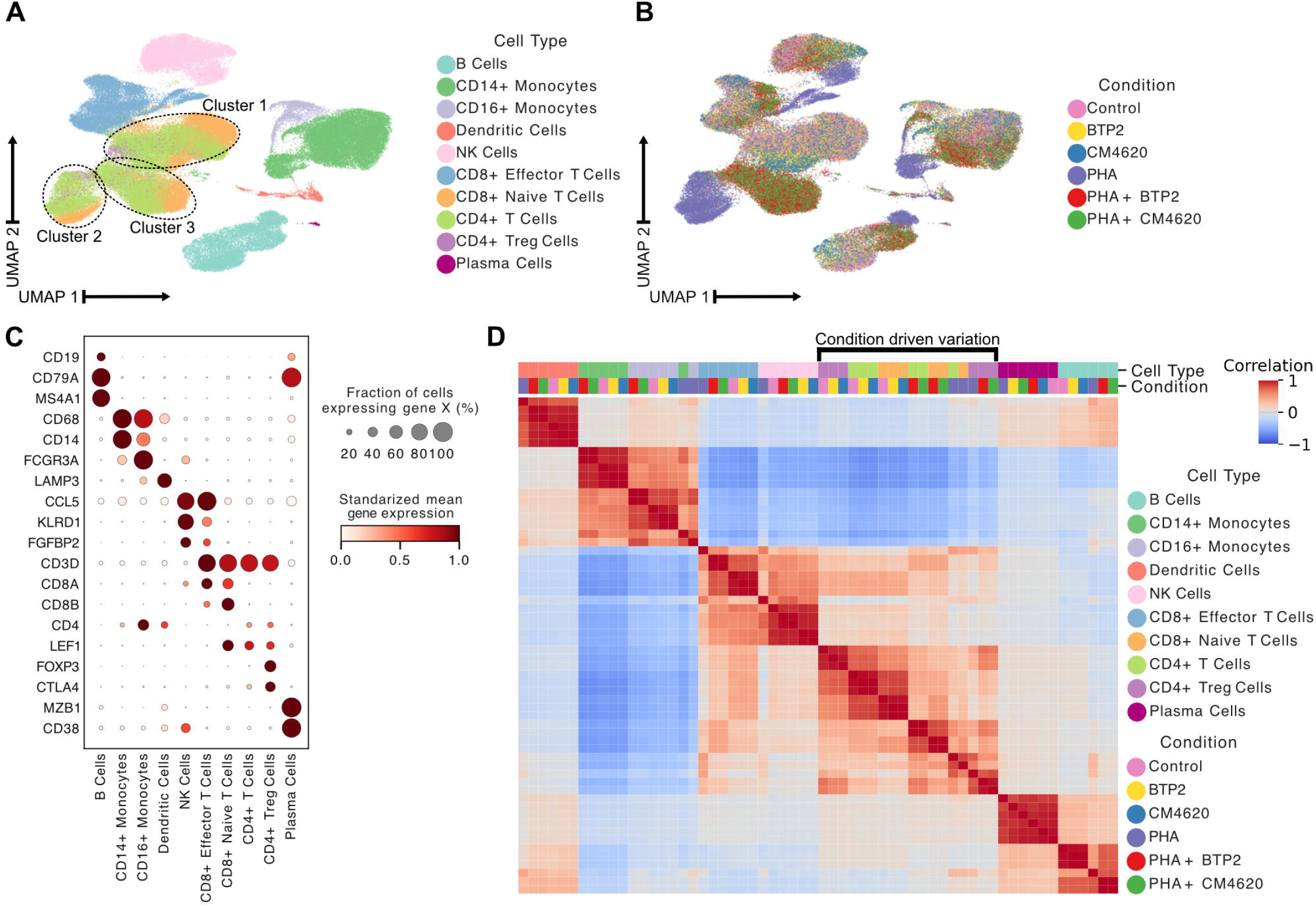
Single-cell RNA sequencing of PHA-stimulated human PBMCs treated with SOCE blockers. **A)** Uniform Manifold Approximation and Projection (UMAP) plot of single-cell transcriptomes of PBMCs from all treatment conditions clustered using gene expression and colored by cell type. **B)** Single-cell transcriptomes of PBMCs from all treatment conditions clustered using gene expression and colored by treatment conditions. **C)** Dot plot showing normalized expression of canonical cell type markers across all cell types. **D)** Heatmap showing pairwise Pearson correlation of single-cell transcriptomes of all cell types split by condition.

### BTP2 and CM4620 induce nearly identical transcriptional effects in human PBMCs

To investigate if the two SOCE blockers have similar effect on the transcriptomes of different immune cell types, we calculated the cosine similarity score and Euclidean distance between pairs of conditions for each cell type (**Methods**). We used cosine similarity as a metric for similarity in gene programs that are impacted and Euclidean distance as a measure of similarity in the magnitude of gene expression. The unstimulated controls had the highest cosine similarity scores and lowest Euclidean distances across all cell types (**Fig. 3**). This is consistent with the observation that the unstimulated controls cluster together on the embedded space. As expected, we observed significantly lower cosine similarity and a higher Euclidean distance when comparing PHA-stimulated samples to controls or PHA-stimulated BTP2 or CM4620 treated samples to PHA-stimulated untreated samples (**Fig. 3**). Interestingly, we observe that the two stimulated drug conditions also demonstrated high similarity based on both metrics, suggesting that the treatment with BTP2 or CM4620 elicited similar transcriptional changes both in terms of gene programs that were affected but also the magnitude of change in the expression of these genes. To further check if there are any differences between the two drugs, we performed a direct DGEA between unstimulated BTP2 treated cells and unstimulated CM4620 treated cells for each cell type, as well as between the PHA-stimulated cells treated with the two drugs for each cell type. This analysis revealed minimal gene expression differences between the cells treated with the two drugs, BTP2 and CM4620, irrespective of their activation state (**Supp. Fig. 10A & 10B, Supp. Table 7-8**). Overall, we find that despite targeting different molecules in the CRAC pathway, SOCE blockade with BTP2 and CM4620 result in an almost identical transcriptional effect on human PBMCs.

**Figure 3:**
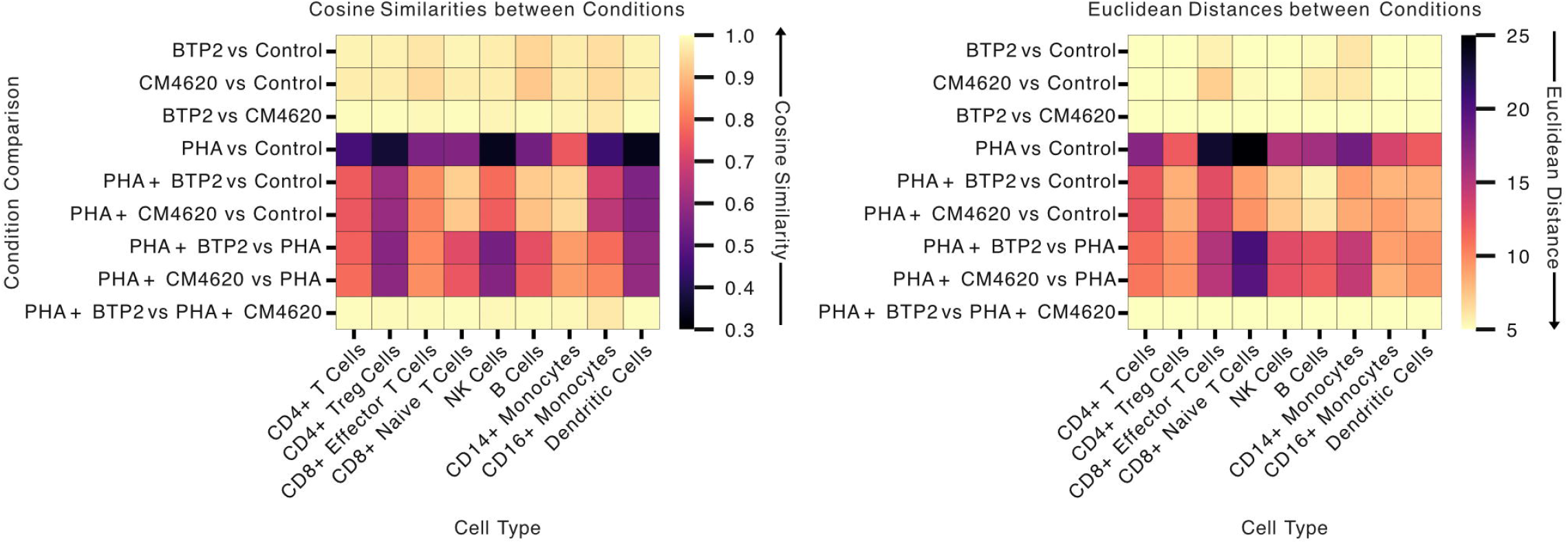
Similarity metrics and DGE analysis on cells treated with the two SOCE blockers: BTP2 and CM4620. Heatmaps show the Euclidean distance and cosine similarity between pairs of treatment conditions for each cell type.

### SOCE blockers suppress inflammation without affecting the anti-inflammatory programs

Given the impact of SOCE blockade on transcription factors crucial for T cell activation and cytokine release, we next characterized, at the single cell level, how BTP2 and CM4620 influence the inflammation-associated molecular pathways in PHA-stimulated T cells. To first characterize the phenotype of PHA-stimulated cytotoxic T cells, we compared PHA-stimulated T cells to unstimulated controls. DGEA revealed a significant upregulation of 1752 genes and a significant downregulation of 188 genes in PHA-stimulated CD8^+^ Effector T cells and upregulation of 929 genes and a significant downregulation of 178 genes in PHA-stimulated NK cells (log_2_ fold-change > 1.0 and adjusted p-value < 0.01, **Supp. Fig. 11A & 11B, Supp. Table 9-10**). Gene ontology analysis of the most significantly upregulated (log_2_ fold-change > 1.0 and adjusted p-value < 0.01) genes in CD8^+^ Effector T cells revealed an enrichment of pathways such as leukocyte mediated cytotoxicity and T cell mediated cytotoxicity (**Supp. Fig. 11C & 11D**). In NK cells, gene set enrichment analysis of upregulated (log_2_ fold-change > 1.0 and adjusted p-value < 0.01) genes revealed an enrichment of ontology terms such as leukocyte mediated cytotoxicity and regulation of leukocyte mediated cytotoxicity (**Supp. Fig. 11C & 11D**). We then generated gene programs using a curated list of 18 genes found in T cell cytotoxicity associated gene ontology terms and used it to estimate cytotoxicity module score for all cells across four conditions (**Methods, Fig. 4A, Supp. Fig. 12A & 12B**). Comparison of cytotoxicity scores for NK cells and all T cell types across four treatment conditions showed that for NK cells and effector CD8^+^ cytotoxic T cells cytotoxicity score was found to increase in PHA-stimulated cells (**Fig. 4B**). As expected, the cytotoxicity score decreased in PHA-stimulated cells treated with BTP2 or CM4620 as compared to PHA-stimulated cells left untreated, which is also consistent with what we have seen using bulk RNA-seq previously (**Fig. 4B**). Interestingly, the cytotoxicity score for both these cell types dropped to levels similar to the levels in unstimulated controls. We calculated Cohen’s distance for all conditions when compared to unstimulated control and found that for NK cells and effector CD8^+^ T cells, Cohen’s distance was maximum for PHA-stimulated cells and the score subsequently decreased for PHA-stimulated cells treated with BTP2 or CM4620 (**Fig. 4C**). Visualizing the expression GZMB and PRF1 specifically, their expression is increased in PHA stimulated PBMCs followed by a decrease after drug treatment in both CD8^+^ effector T cells and NK cells (Fig. 4D). This is consistent with both the qPCR and bulk RNA-seq results (**Supp. Fig 3F & 4**).

**Figure 4:**
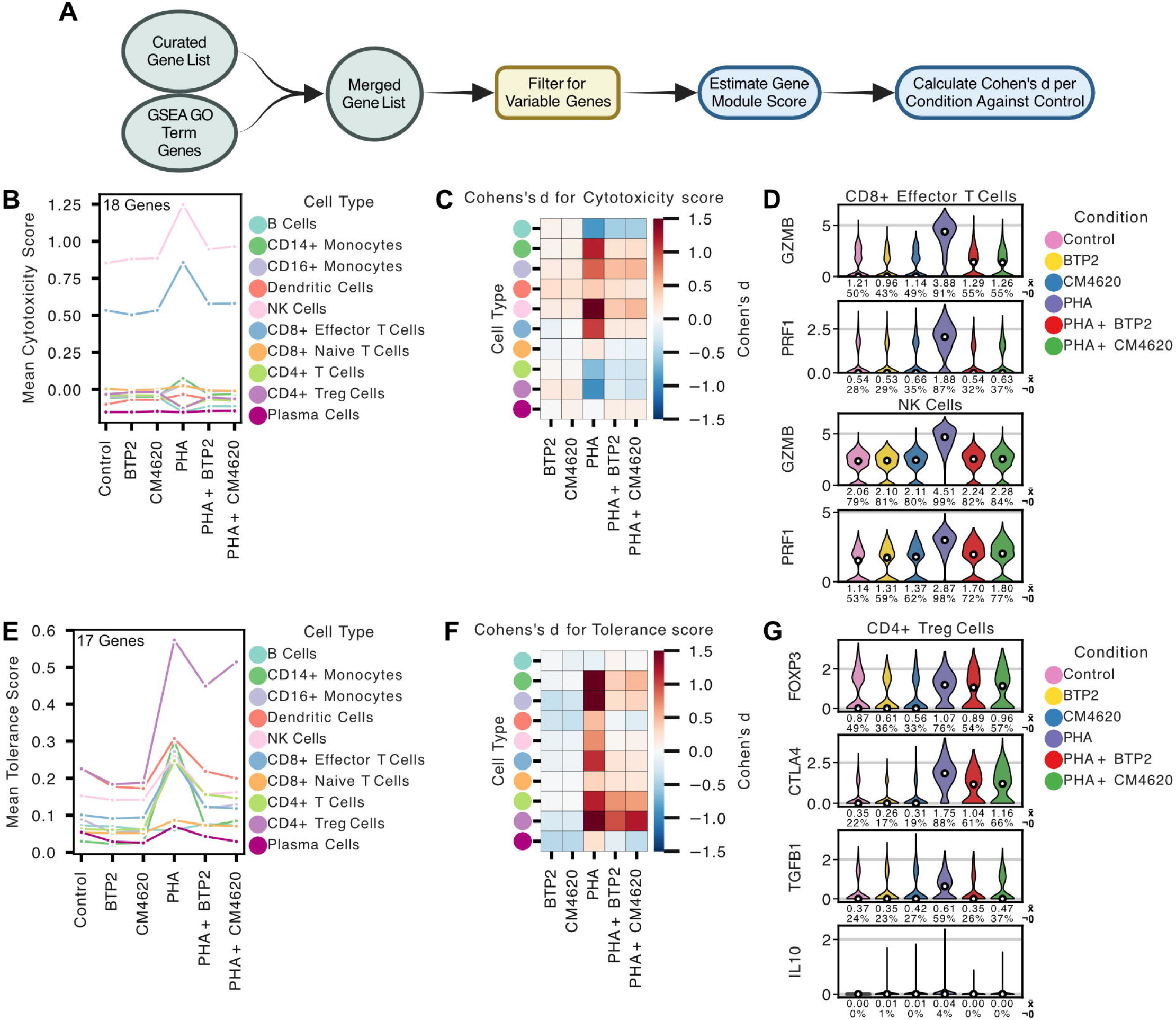
Comparison of cytotoxic and tolerance gene expression in PHA-stimulated PBMCs treated with SOCE blockers. **A)** Flowchart overview of analysis for estimating gene module scores and Cohen’s dissimilarity metric between conditions. **B)** Line plot showing the mean cytotoxicity scores, generated using 18 cytotoxicity genes, for each cell type split across treatment conditions. **C)** Heatmap of Cohen’s d metric for cytotoxicity for each cell type comparing each condition against the PBMCs (Control) condition. **D)** Violin plots showing log transformed normalized expression of GZMB and PRF1 in CD8^+^ Effector T cells and NK cells split by the six condition groups. Median is indicated by the white dots, the average by the x□ row, and the percentage of cells that have a non-zero value by the ¬0 row. **E)** Line plot showing the mean tolerance scores, generated using 17 tolerance genes, for each cell type split across treatment conditions. **F)** Heatmap of Cohen’s d metric for tolerance for each cell type comparing each condition against the PBMCs (Control) condition. **G)** Violin plots showing log transformed normalized expression of FOXP3, CTLA4, TGFB1, and IL10 in CD4^+^ Treg cells split by the six condition groups. Median is indicated by the white dots, the average by the x□ row, and the percentage of cells that have a non-zero value by the ¬0 row.

Next, we generated a tolerance/anti-inflammatory gene module using a curated list of 17 genes in ontology terms associated with tolerance and anti-inflammatory pathways (**Fig. 4A, Supp. Fig. 12C & 12D**). We then compared tolerance gene module scores across all four conditions for all immune cell subtypes. Except for CD8^+^ Naive T cells, B cells, and plasma cells, stimulation with PHA resulted in an increase of anti-inflammatory/tolerance gene expression as demonstrated by an increase in tolerance module score (**Fig. 4E**). However, CD4^+^ T-reg cells were the only stimulated cell type that maintained a high tolerance score even after treatment with either of the two drugs BTP2 or CM4620. All other immune cells demonstrated a drop in tolerance score after drug treatment (**Fig. 4E**). We then calculated Cohen’s d metric for each condition compared to the unstimulated untreated condition for CD4^+^ T-reg cells and found that the Cohen’s d metric was large not only for the PHA stimulated cells, but also for the stimulated cells treated with BTP2 or CM4620 (**Fig. 4F**). Consistent with the results of the qPCRs, the expression of FOXP3 and CTLA4 is increased in stimulated PBMCs and maintained at a level higher than the controls even after drug treatment in CD4^+^ Treg cells (Fig. 4G). Furthermore, we performed DGEA for PHA-stimulated CD4^+^ T-reg cells treated with BTP2 or CM4620 compared to unstimulated control (**Supp. Table 11-12**). The DGEA analysis revealed upregulated (log_2_ fold-change > 1.0 and adjusted p-value < 0.01) genes associated with ontology terms such as regulation of T cell activation and negative regulation of T cell receptor signaling pathway for both drugs (**Supp. Fig. 13**). The GO term for regulation of T cell activation in the PHA-stimulated CM4620-treated PBMCs showed an enrichment in 26 genes: CORO1A, XBP1, PRELID1, PRNP, SIRPG, IL2RG, HSPD1, ICOS, TNFRSF9, CD74, CTLA4, ACTB, HSPH1, BATF, LRRC32, PNP, IL2RA, CD28, TNFRSF1B, CBFB, SIT1, ACTL6A, PTPN6, SELENOK, LGALS1, and RAC2, many of which are known to be associated with T-reg cell tolerance. Similarly, for the same GO term, PHA-stimulated BTP2-treated cells showed an enrichment in genes such as CORO1A, XBP1, PRELID1, PRNP, IL2RG, HSPD1, TIGIT, TNFRSF9, CTLA4, ACTB, HSPH1, BATF, LRRC32, PNP, CD28, PTPN6, SOCS1, SELENOK, LGALS1, and RAC2, many of which are associated with T-reg cell tolerance. Overall, we report that regulatory T cell mediated tolerance associated genes are maintained in PHA-stimulated PBMCs following treatment with SOCE blockers unlike the cytotoxic gene set down regulated by the SOCE blockers.

## DISCUSSION

Blocking calcium influx via store-operated calcium entry (SOCE) impairs various immune cell functions, including cytokine production, proliferation, and cytotoxic activity. Genetic deletion or pharmacological inhibition of SOCE components can impair T cell activation and function^30^. However, complete SOCE inhibition may cause immunodeficiency, as seen in patients with CRAC channel mutations^31^. Therefore, partial or selective inhibition presents a viable therapeutic approach. Our findings show that SOCE blockade with BTP2 or CM4620 suppresses cytotoxic gene expression while sparing activation-induced anti-inflammatory gene expression. Specifically, in PHA-stimulated PBMCs, cytotoxicity-associated genes in CD8+ effector T cells and NK cells dropped to levels comparable to unstimulated PBMCs, while CD4+ T-reg cells maintained their tolerance phenotype. These findings support our hypothesis that SOCE blockers selectively modulate immune responses, shifting the balance toward tolerance and away from inflammation. This selective suppression could provide a promising approach for immune suppression in autoimmune diseases and transplantation.

Current immunosuppressive therapies in transplantation rely on broad-spectrum agents such as calcineurin inhibitors (e.g., cyclosporine, tacrolimus), mTOR inhibitors (e.g., sirolimus), corticosteroids, and antimetabolites like mycophenolate mofetil. While these drugs improve graft survival, they cause significant side effects, including nephrotoxicity, infections, malignancies, and metabolic complications^32^. Moreover, they indiscriminately suppress the immune system, affecting both pathogenic and protective immune responses. Our findings align with ongoing efforts to develop selective immunosuppressive strategies that minimize these adverse effects. The ability to reduce cytotoxicity in effector cells while preserving regulatory mechanisms could enhance graft survival and function. By maintaining Treg activity, SOCE blockade may promote immune tolerance, reducing chronic rejection— a major cause of long-term graft loss. Additionally, preserving anti-inflammatory responses may aid tissue repair and reduce graft inflammation^33^. This dual effect of attenuating harmful immune responses while supporting regulatory and reparative mechanisms addresses a critical need in transplantation medicine. By specifically targeting SOCE we can attenuate harmful immune responses while preserving beneficial ones.

Further studies in transplantation models are needed to evaluate SOCE blockers’ efficacy in preventing acute and chronic rejection, assessing immunological outcomes, pharmacokinetics, optimal dosing, and potential toxicity. Long-term effects on immunity, including infection risk, malignancies, and vaccine responses, should also be investigated. Despite these open questions, our study contributes to the evolving landscape of transplant immunology by presenting SOCE blockade as a targeted immunosuppressive strategy. By selectively inhibiting cytotoxic immune responses while preserving regulatory and anti-inflammatory functions, SOCE blockers have the potential to improve transplantation outcomes significantly.

## ACKNOWLEDGEMENTS

We thank the members of the Genomics Resources Core Facility of the Weill Cornell Medical College, New York, New York, USA for the RNA sequencing studies. M.S. was supported by an award (R37 NIH MERIT AI051652) from the National Institute of Allergy and Infectious Diseases, National Institutes of Health. MM was supported by R01AI151059 (to I.D.V.) and R21AI164093 (to I.D.V.). The statements made herein are solely the responsibility of the authors.

## AUTHOR CONTRIBUTIONS

All authors contributed to the study design and experimental design. D.S., C.L., T.M. and M.S., contributed to the experiments. A.S., M.M., and D.S. contributed to data analysis. A.S., M.M., D.S., C.L., I.D.V., T.M., and M.S. contributed to the writing of the manuscript. All authors provided feedback and approved the final version of the manuscript.

## DATA AVAILABILITY

The authors declare that all sequencing data supporting the findings of this study have been deposited in NCBI’s Gene Expression Omnibus (GEO) with GEO series accession number GSE295784. All other data supporting the findings in this study are included in the main article and associated files.

## CODE AVAILABILITY

All code needed to reproduce the RNA-seq analysis presented in this study have been deposited on the GitHub repository: https://github.com/AndreasStephanou/SOCE_Blockade).

## REFERENCES

1. Dolmetsch, R. E., Xu, K. & Lewis, R. S. Calcium oscillations increase the efficiency and specificity of gene expression. Nature 392, 933–936 (1998).

2. Berridge, M. J. & Irvine, R. F. Inositol phosphates and cell signalling. Nature 341, 197–205 (1989).

3. Prakriya, M. & Lewis, R. S. Store-Operated Calcium Channels. Physiol. Rev. 95, 1383–1436 (2015).

4. Hogan, P. G., Lewis, R. S. & Rao, A. Molecular basis of calcium signaling in lymphocytes: STIM and ORAI. Annu. Rev. Immunol. 28, 491–533 (2010).

5. Chitwood, K. K. & Heim-Duthoy, K. L. Immunosuppressive Properties of Calcium Channel Blockers. Pharmacother. J. Hum. Pharmacol. Drug Ther. 13, 447–454 (1993).

6. Weir, M. R. Therapeutic benefits of calcium channel blockers in cyclosporine-treated organ transplant recipients: Blood pressure control and immunosuppression. Am. J. Med. 90, S32–S36 (1991).

7. Nieto-Felipe, J. et al. Role of Orai-family channels in the activation and regulation of transcriptional activity. J. Cell. Physiol. 238, 714–726 (2023).

8. Trebak, M. & Kinet, J.-P. Calcium signalling in T cells. Nat. Rev. Immunol. 19, 154–169 (2019).

9. Srikanth, S. & Gwack, Y. Orai1-NFAT Signalling Pathway Triggered by T Cell Receptor Stimulation. Mol. Cells 35, 182–194 (2013).

10. Hackam, D. J., Rotstein, O. D., Schreiber, A., Zhang, W. & Grinstein, S. Rho is Required for the Initiation of Calcium Signaling and Phagocytosis by Fcγ Receptors in Macrophages. J. Exp. Med. 186, 955–966 (1997).

11. Waldron, R. T. et al. The Orai Ca2+ channel inhibitor CM4620 targets both parenchymal and immune cells to reduce inflammation in experimental acute pancreatitis. J. Physiol. 597, 3085–3105 (2019).

12. Lewis, S. et al. Combination of the CRAC Channel Inhibitor CM4620 and Galactose as a Potential Therapy for Acute Pancreatitis. Function 5, zqae017 (2024).

13. Bruen, C. et al. Auxora for the Treatment of Patients With Acute Pancreatitis and Accompanying Systemic Inflammatory Response Syndrome: Clinical Development of a Calcium Release-Activated Calcium Channel Inhibitor. Pancreas 50, 537–543 (2021).

14. Ohga, K. et al. The suppressive effects of YM-58483/BTP-2, a store-operated Ca2+ entry blocker, on inflammatory mediator release in vitro and airway responses in vivo. Pulm. Pharmacol. Ther. 21, 360–369 (2008).

15. Ohga, K., Takezawa, R., Arakida, Y., Shimizu, Y. & Ishikawa, J. Characterization of YM-58483/BTP2, a novel store-operated Ca2+ entry blocker, on T cell-mediated immune responses in vivo. Int. Immunopharmacol. 8, 1787–1792 (2008).

16. Yoshino, T. et al. YM-58483, a selective CRAC channel inhibitor, prevents antigen-induced airway eosinophilia and late phase asthmatic responses via Th2 cytokine inhibition in animal models. Eur. J. Pharmacol. 560, 225–233 (2007).

17. Shankaranarayanan, D. et al. Blockade of store-operated calcium entry by BTP2 preserves anti-inflammatory gene expression in human peripheral blood mononuclear cells. Hum. Immunol. 85, 111144 (2024).

18. Mantri, M. et al. Spatiotemporal single-cell RNA sequencing of developing chicken hearts identifies interplay between cellular differentiation and morphogenesis. Nat. Commun. 12, 1771 (2021).

19. Mantri, M. et al. Spatiotemporal transcriptomics reveals pathogenesis of viral myocarditis. Nat. Cardiovasc. Res. 1, 946–960 (2022).

20. Dobin, A. et al. STAR: ultrafast universal RNA-seq aligner. Bioinforma. Oxf. Engl. 29, 15–21 (2013).

21. Love, M. I., Huber, W. & Anders, S. Moderated estimation of fold change and dispersion for RNA-seq data with DESeq2. Genome Biol. 15, 550 (2014).

22. Huber, W., von Heydebreck, A., Sültmann, H., Poustka, A. & Vingron, M. Variance stabilization applied to microarray data calibration and to the quantification of differential expression. Bioinforma. Oxf. Engl. 18 Suppl 1, S96–104 (2002).

23. Tibshirani, R. Estimating Transformations for Regression via Additivity and Variance Stabilization. J. Am. Stat. Assoc. 83, 394–405 (1988).

24. Zhu, A., Ibrahim, J. G. & Love, M. I. Heavy-tailed prior distributions for sequence count data: removing the noise and preserving large differences. Bioinformatics 35, 2084–2092 (2019).

25. Wolf, F. A., Angerer, P. & Theis, F. J. SCANPY: large-scale single-cell gene expression data analysis. Genome Biol. 19, 15 (2018).

26. Subramanian, A. et al. Gene set enrichment analysis: A knowledge-based approach for interpreting genome-wide expression profiles. Proc. Natl. Acad. Sci. 102, 15545–15550 (2005).

27. Fang, Z., Liu, X. & Peltz, G. GSEApy: a comprehensive package for performing gene set enrichment analysis in Python. Bioinformatics 39, btac757 (2023).

28. Ashburner, M. et al. Gene Ontology: tool for the unification of biology. Nat. Genet. 25, 25–29 (2000).

29. The Gene Ontology Consortium et al. The Gene Ontology knowledgebase in 2023. Genetics 224, iyad031 (2023).

30. Gwack, Y. et al. Hair Loss and Defective T- and B-Cell Function in Mice Lacking ORAI1. Mol. Cell. Biol. 28, 5209 (2008).

31. Feske, S., Picard, C. & Fischer, A. Immunodeficiency due to mutations in ORAI1 and STIM1. Clin. Immunol. Orlando Fla 135, 169–182 (2010).

32. Ekberg, H. et al. Reduced Exposure to Calcineurin Inhibitors in Renal Transplantation. N. Engl. J. Med. 357, 2562–2575 (2007).

33. Lechler, R. I., Sykes, M., Thomson, A. W. & Turka, L. A. Organ transplantation--how much of the promise has been realized? Nat. Med. 11, 605–613 (2005).

