## Supplemental Methods and Figures for "Single-cell transcriptomics reveals targeted modulation of inflammatory repertoire by SOCE blockers"

### PBMC Isolation and Stimulation

PBMCs were isolated from peripheral blood of healthy volunteers using Ficoll-Paque™ density gradient centrifugation. PBMCs ($1\times10^6$ cells/mL) were incubated under the following conditions: control (without PHA or inhibitors), 1000 nM BTP2, 2 µg/mL PHA, 2 µg/mL PHA + 1000 nM BTP2, or 2 µg/mL PHA + 1000 nM CM4620 for 16 hours at 37°C in 95% air and 5% $CO_2$. DMSO was used as the solvent control. After incubation, PBMCs were collected for downstream RNA isolation and sequencing.

### RNA Isolation and Quantitative PCR

PBMCs were washed with 1 mL PBS and centrifuged at 13,400 × g for 2 minutes at room temperature. After discarding the supernatant, cells were lysed in 50 µL RNAlater and 350 µL Buffer RLT containing β-mercaptoethanol. RNA was extracted using the RNeasy Mini Kit (Qiagen) according to the manufacturer's protocol. RNA purity (A260/A280 ratio) and quantity were assessed using a NanoDrop® One UV-Vis spectrophotometer (Thermo Scientific).

For RT-qPCR, RNA was reverse transcribed to cDNA, and absolute mRNA copy numbers were quantified using preamplification-enhanced RT-qPCR assays. Primer sequences and gene-specific TaqMan probes were previously published (reference 17). Four independent experiments were performed using PBMCs from four healthy donors.

### Bulk RNA Library Preparation and Sequencing

RNA samples were treated with RNase-Free DNase (Qiagen) to remove residual DNA. RNA library preparation was conducted using the TruSeq RNA Library Prep Kit (Illumina), and sequencing was performed at Weill Cornell Medicine using the NovaSeq 6000 system with 100 × 2 paired-end reads.

### Single-Cell RNA Sequencing (scRNA-seq)

Live PBMCs were counted using a TC20 cell counter (Bio-Rad), and approximately 2 million cells per sample were labeled using the 3' CellPlex Kit Set A (10X Genomics). Cell suspensions were combined to generate gene expression libraries using Chromium Single Cell 3' Reagent Kits v3.1 with Feature Barcoding technology. Libraries were sequenced on a NovaSeq 6000 (Illumina).

### Bulk RNA-seq Data Processing

Raw sequencing reads were aligned to the human genome (GRCh38) using STAR (v-2.7.10b). Gene expression counts were processed with DESeq2 (v-1.44.0), where genes with fewer than ten copies across all samples were removed. Variance-stabilizing transformation (VST) was used for normalization, and batch correction was applied using limma (v-3.60.0). Principal Component Analysis (PCA) and Differential Gene Expression Analysis (DGEA) were performed using DESeq2's Wald test, with effect size shrinkage applied using apeglm (v-1.26.1).

### scRNA-seq Preprocessing and Analysis

Raw gene expression matrices were processed using Scanpy (v.1.10.1). Cells with fewer than 500 genes, fewer than 1,000 UMI counts, or mitochondrial content exceeding 20% were removed. Doublets were detected using Scrublet (v.0.2.3) and excluded. Data were normalized to 10,000 counts per cell and log-transformed. Highly variable genes were identified using min_mean = 0.1 and min_dispersion = 0.2.

PCA was performed using the top 20 principal components, followed by Leiden clustering (resolution = 0.3) and UMAP embedding. Clusters were annotated based on canonical immune cell markers.

For T cell subpopulation analysis, CD4+ and CD8+ T cells were subsetted and reanalyzed using 12 PCs and Leiden clustering (resolution = 2.0). CD8+ T cells were further classified as effector or naïve based on cytotoxic marker expression. CD4+ regulatory T cells (T-regs) were identified using FOXP3 expression with Leiden clustering (resolution = 2.5).

**Cell Type Correlations and Similarity Metrics**

PBMC cell types were split by condition into 60 groups, and mean gene expression was calculated for each. Pearson correlation coefficients were computed using Pandas (v.2.2.1), and hierarchical clustering was performed using SciPy (v.1.11.4) with average linkage and correlation distance metrics.

To assess similarity between conditions, centroids for each cell type were computed using mean gene expression. Euclidean distances and cosine similarity were calculated using SciPy (v-1.11.4) and scikit-learn (v.1.4.2), respectively. Comparisons included control vs. drug-treated conditions and between different drug treatments.

**Gene Module Scores and Cohen's d Calculation**

Cytotoxicity gene module scores were derived by identifying highly variable genes in CD8+ effector T cells and NK cells, supplemented with GSEA GO term "GOBP Positive Regulation of T Cell Mediated Cytotoxicity" and known markers (NKG7, IFNG, PRF1). Scanpy's score_genes function was used to compute module scores, and Cohen's d was calculated for effect size estimation.

Similarly, tolerance scores were derived from CD4+ T-reg cell genes identified using GO terms for T cell tolerance induction. IL10 was included despite low expression. Tolerance scores and Cohen's d were calculated using the same methodology.

**Differential Gene Expression and Gene Set Enrichment Analysis (GSEA)**

DEGs were identified using the Wilcoxon rank-sum test (Scanpy's rank_genes_groups) with adjusted $p < 0.01$. Upregulated genes were defined as log2 fold-change > 1, and downregulated genes as log2 fold-change < -1. GSEA was performed using GSEApy (v.1.0.3) with the "c5.all.v2023.2.Hs.symbols.gmt" gene set. DGE and GSEA were repeated for control vs. drug-treated conditions across all cell types, with the top 10 GO terms visualized.

**SUPPLEMENTAL FIGURES**

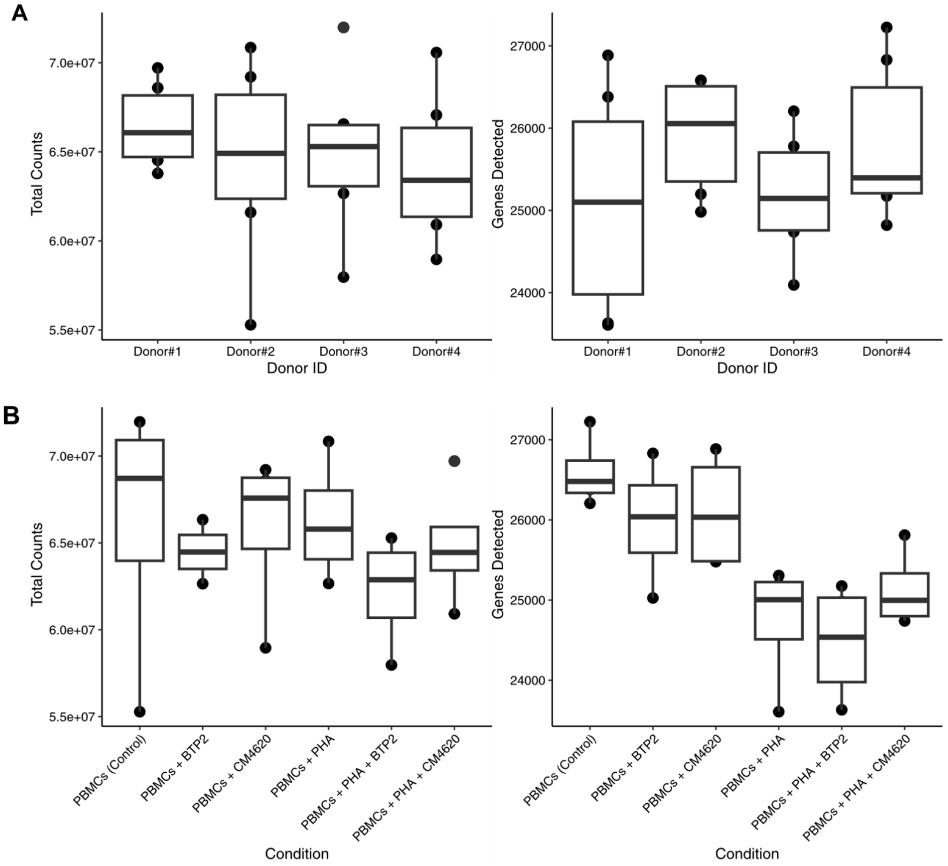


**Supplementary Figure 1**: **Quality control for bulk RNA-seq samples across different donors and treatment conditions. A)** Box plot showing total counts and genes detected in individual samples across donors (#1 to #4). **B)** Box plot showing total counts and genes detected in individual samples across different treatment conditions: PBMCs (Control), PBMCs treated with BTP2 or CM4620, PBMCs stimulated with PHA, and PHA-stimulated PBMCs treated with BTP2 or CM4620. Each dot represents an individual sample.

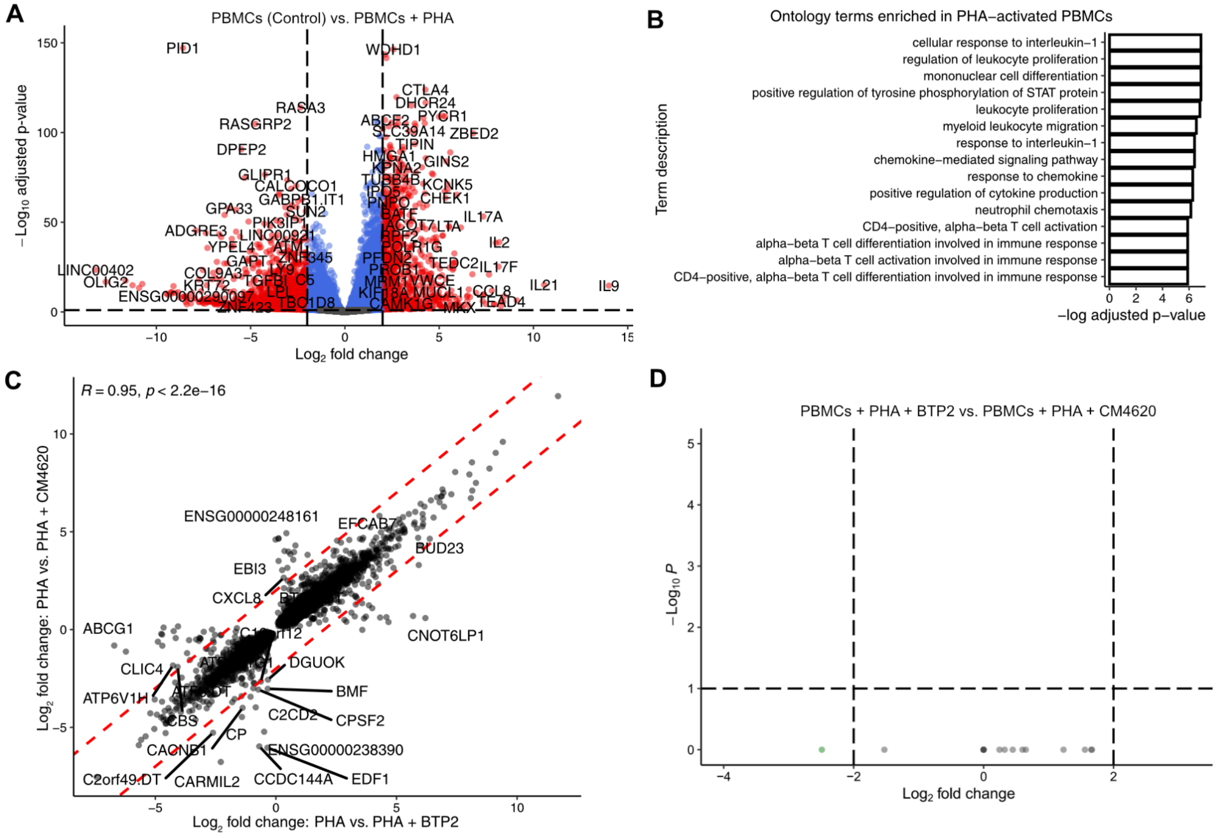


**Supplementary Figure 2**: Differential gene expression analysis under various conditions. **A)** Volcano plot showing the $\log_2$ fold-change and $-\log_{10}$ adjusted p-value for genes when compared between PBMCs (Control) and PBMCs stimulated with PHA. The dotted lines define the $\log_2$ fold-change threshold of +/- 2.0 and adjusted p-value threshold of 0.01. **B)** Bar plot showing top 15 ontology terms enriched for genes upregulated in PBMCs + PHA as compared to PBMCs (Control) with $\log_2$ fold-change > 5.0 and adjusted p-value < 0.01. **C)** Scatter plot showing the correlation between log fold-change for genes significantly altered in PBMCs + PHA + BTP2 or altered in PBMCs + PHA + CM4620, as compared to PBMCs + PHA. Only significantly altered genes with adjusted p-value < 0.01 in at least one of the two comparisons were included in the plot. **D)** Volcano plot showing the $\log_2$ fold-change and $-\log_{10}$ adjusted p-value for genes when compared between PBMCs + PHA + BTP2 and PBMCs + PHA + CM4620.

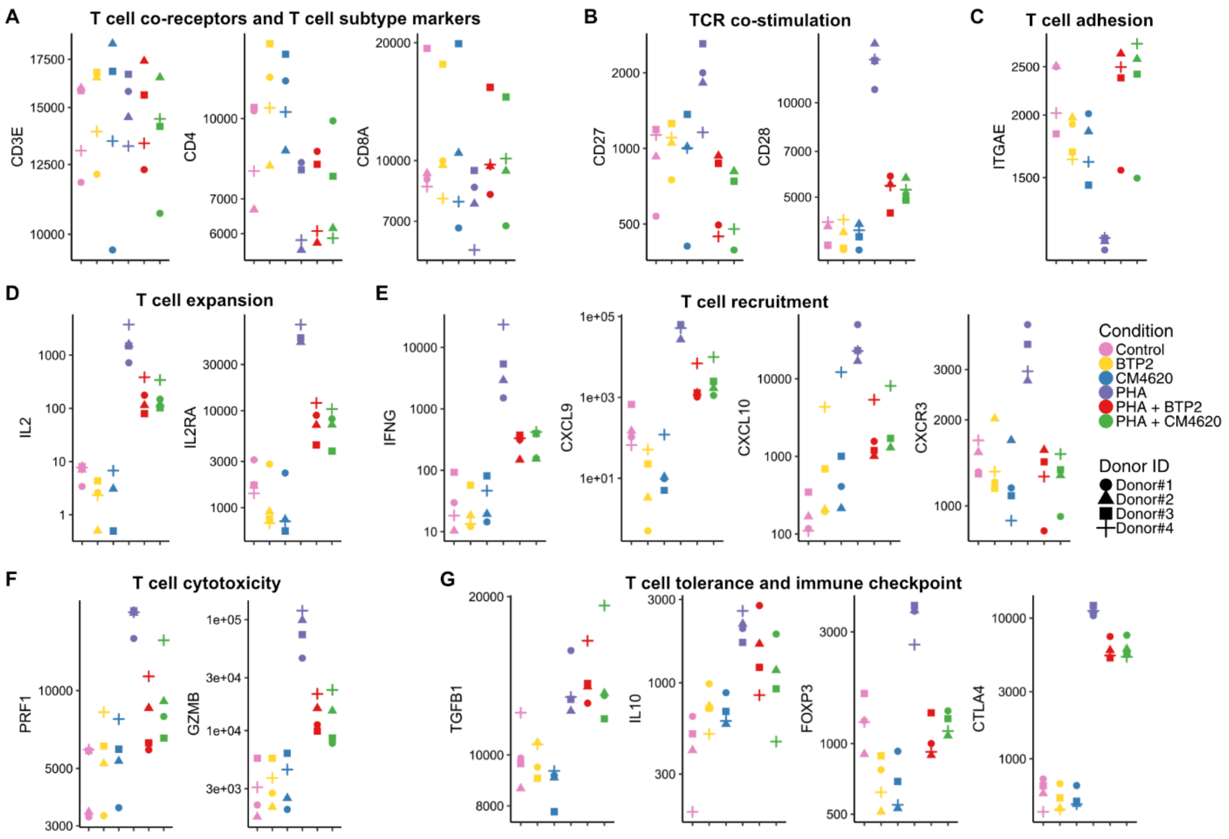


**Supplementary Figure 3: Differential effects of BTP2 and CM4620 on PHA-induced alterations in the expression of mRNAs encoding proteins involved in T cells expansion, recruitment, activation, effector function, and tolerance.** Jitter plots showing the log-normalized counts across the six treatment conditions. **A)** Pan-T cell surface antigen receptor complex CD3E, CD4 expressed on T helper cells and responsible for recognition of antigenic peptide in the context of HLA class II antigens and CD8 expressed on cytotoxic T cells and responsible for recognition of antigenic peptide in the context of HLA class I antigens; **B)** T cell co-stimulation receptors CD27 and CD28; **C)** T cell adhesion marker: surface integrin ITGAE (CD103); **D)** T cell growth factor IL2 and its receptor IL2RA responsible for T cell clonal expansion; **E)** IFNG and IFNG induced T cell recruitment/homing proteins: Chemoattractant molecules and receptors: CXCL9, CXCL10, and CXCR3; **F)** T cell cytotoxic proteins PRF1 and GZMB **G)** Immunosuppressive cytokines TGFB1 and IL10, Treg cell marker FOXP3 and the master negative regulator of immunity CTLA4.

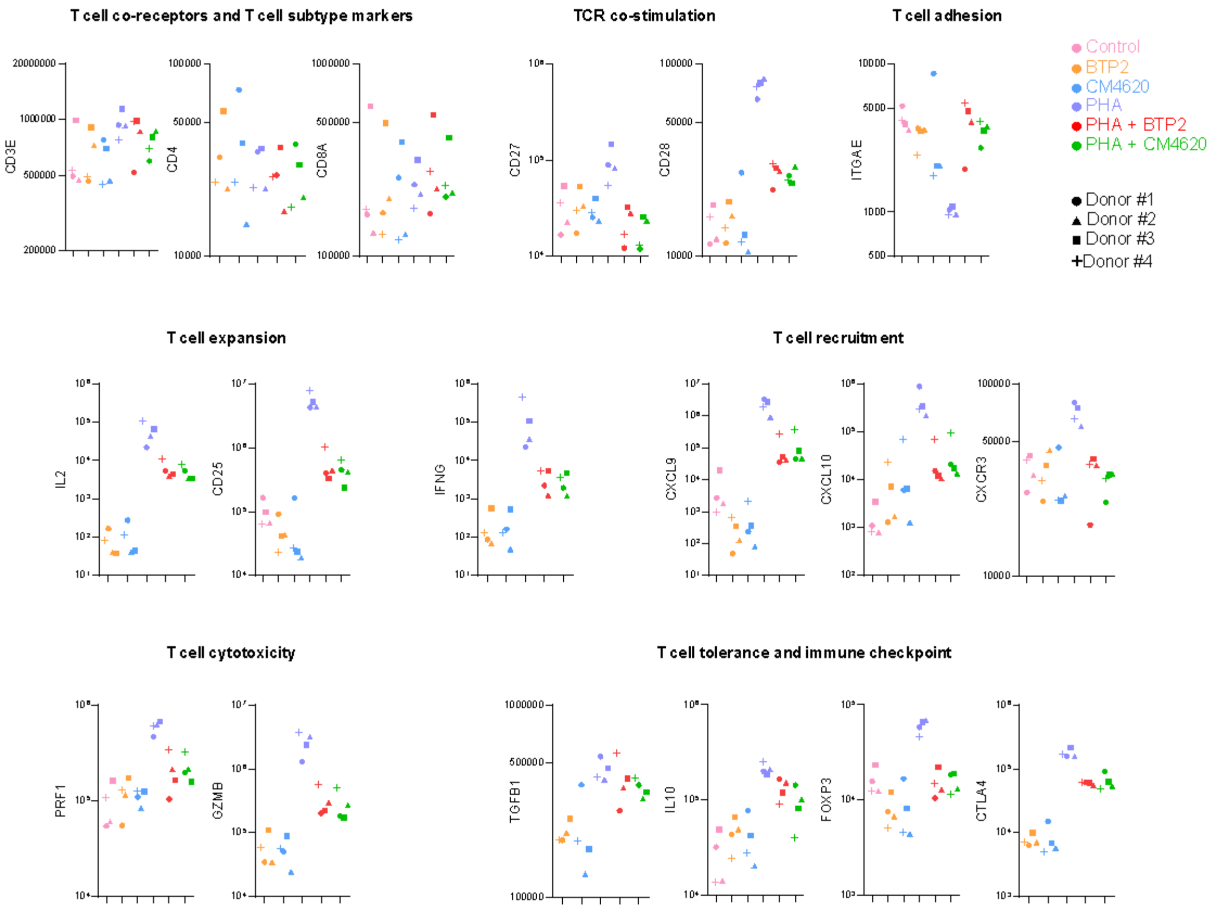


**Supplementary Figure 4: RT-qPCR validation of differential effects of BTP2 and CM4620 on PHA-induced alterations in the expression of mRNAs encoding proteins involved in T cells expansion, recruitment, activation, effector function, and tolerance.** Jitter plots show the log-normalized absolute copy number of mRNA per microgram of total RNA from all six experimental conditions. The expression level of mRNA for T cell receptor CD3E (Kruskal-Wallis P=0.96) and T cell subtypes CD4 (P=0.96) and CD8A (P=0.96) were not altered by either BTP2 or CM4620. ITGAE (CD103) mRNA was significantly downregulated by PHA activation, and this downregulation was significantly reversed by both BTP2 and CM4620. Both BTP-2 and CM4620 significantly and markedly suppressed PHA-induced expression of mRNAs for CD27, CD28, IL-2, CD25, IFNG, CXCL9 and CXCL10, PRF1 (perforin) and GZMB (granzyme B). Neither BTP2 nor CM4620 significantly altered the expression of mRNA encoding immunosuppressive cytokine TGFB1. There was significant but only modest suppression of mRNAs for IL10, FOXP3 and CTLA4. A pair-wise comparison using the Wilcoxon signed-rank test was used to investigate the differences in mRNA copy number between PBMCs+PHA vs. PBMCs+PHA+BTP2 as well as PBMCs+PHA vs. PBMCs+PHA+CM4620. A P-value <0.05 was considered significant for alterations in mRNA copy number.

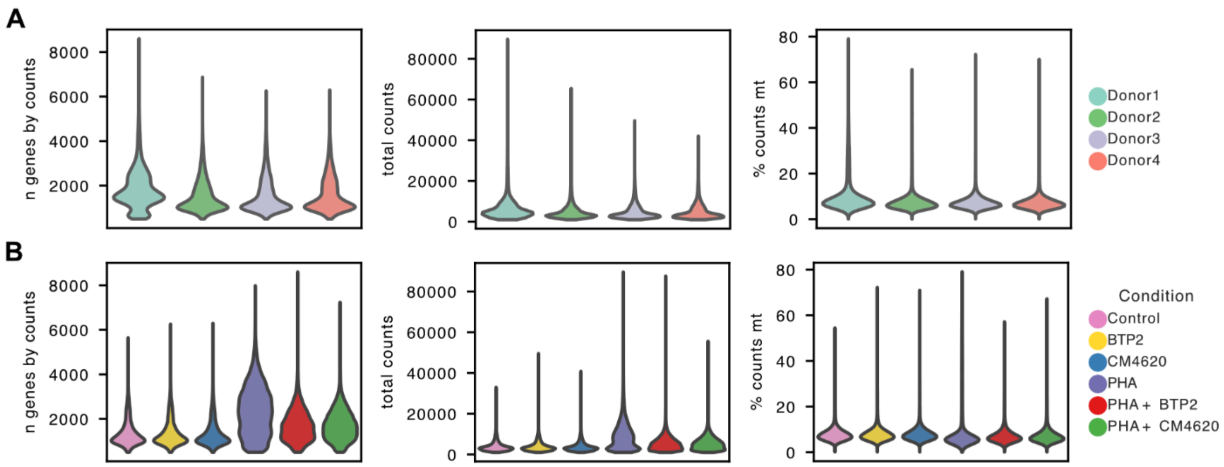


**Supplementary Figure 5: Quality control metrics of scRNA-seq analysis. A-B)** Violin plots showing the number of genes detected per cell (left), total transcript counts per cell (middle), and percentage of mitochondrial transcripts (right) in cells grouped by: **A)** Donor and **B)** Condition. After quality control filtering, we detected a mean of 3,471 to 5,222 transcripts across donors. A higher number of distinct genes as well as a greater total number of transcripts were detected in PBMCs activated with PHA as compared to un-activated PBMCs as well as activated PBMCs treated with BTP2 and CM4620. Interestingly, PHA-activated PBMCs treated with BTP2 or CM4620 exhibited lower transcript diversity and transcript counts compared to PHA-activated cells but still displayed higher values than control PBMCs.

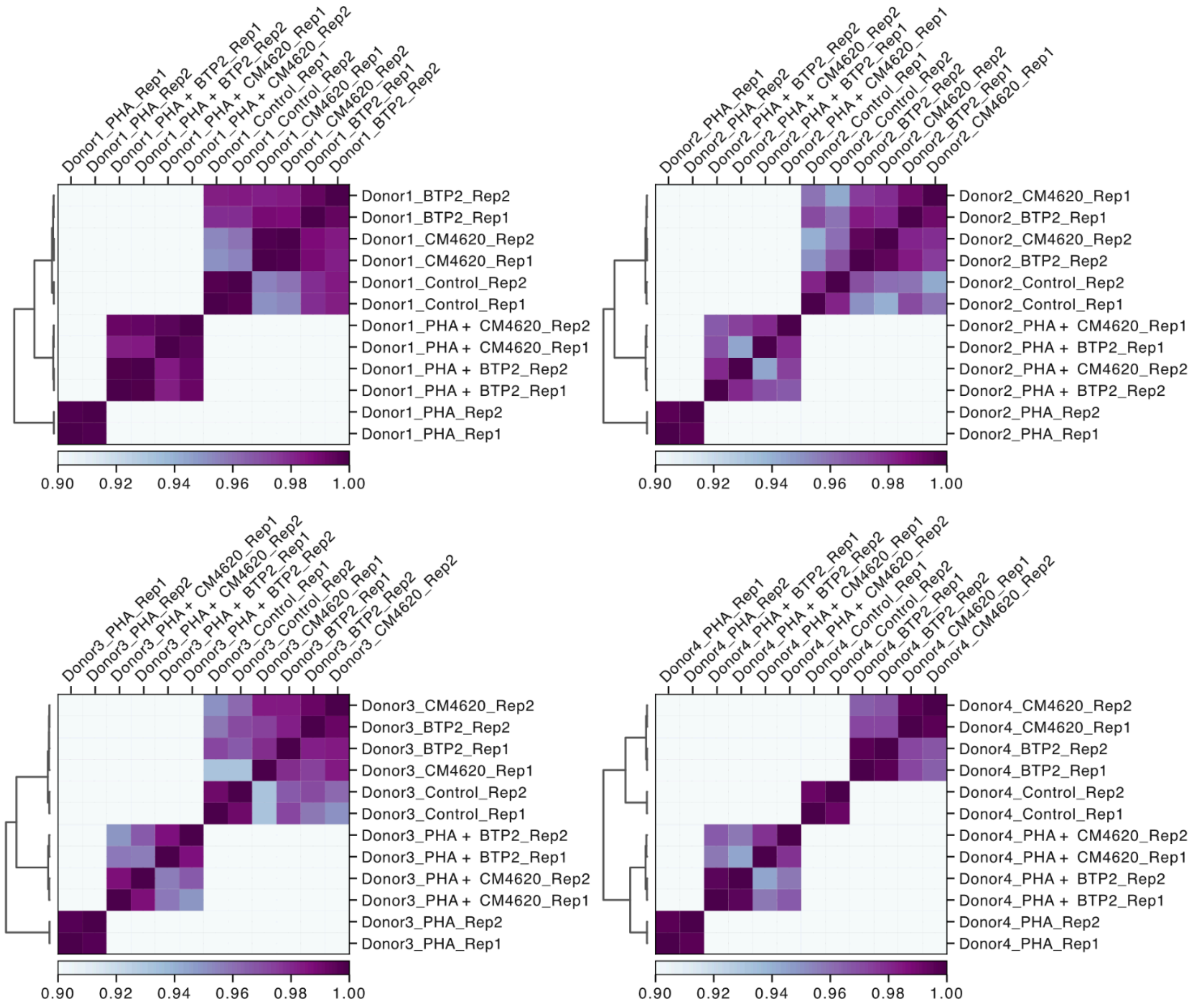


**Supplementary Figure 6**: Heatmaps showing the pairwise Pearson correlation values for all sample replicates across all treatment conditions within a donor. Individual heat maps represent individual donors. Correlation analysis across samples within each donor revealed high correlations, typically ranging from 0.92 to 1.0, between technical replicates indicating strong reproducibility within each condition.

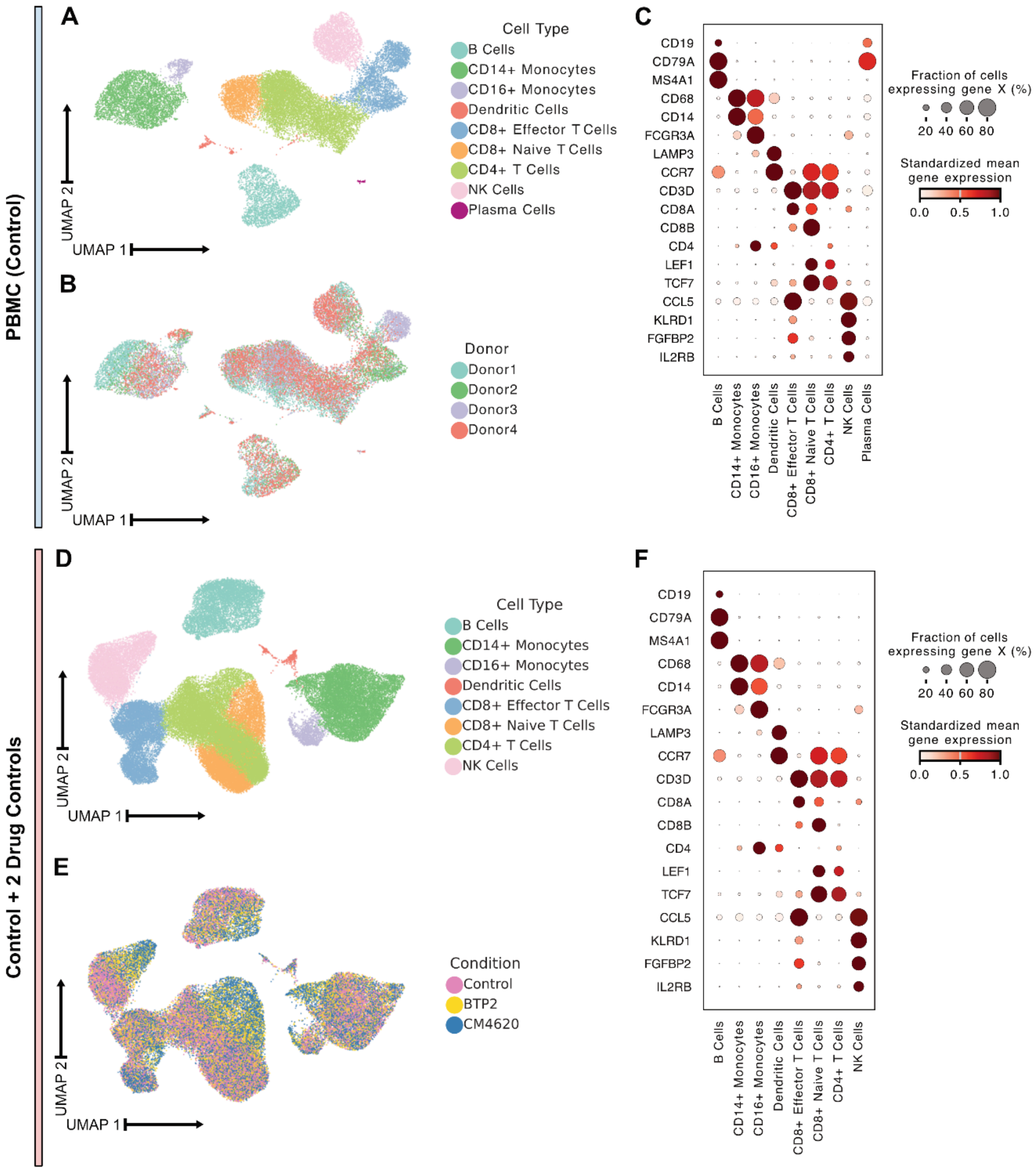


**Supplementary Figure 7: scRNA-seq analysis of just control PBMCs and of control PBMCs and treated but unstimulated PBMCs combined. A)** UMAP plot of single-cell transcriptomes from control PBMCs clustered by gene expression colored by cell type. **B)** UMAP plot of single-cell transcriptomes from control PBMCs clustered by gene expression colored by donor of origin. **C)** Dot plot showing the normalized gene expression for canonical cell type markers in individual cell types detected within control PBMCs. **D)** UMAP plot of single-cell transcriptomes from control PBMCs and unstimulated PBMCs treated with SOCE blockers clustered by gene expression colored by cell type. **E)** UMAP plot of single-cell transcriptomes from control PBMCs and unstimulated PBMCs treated with SOCE blockers clustered by gene expression colored by donor of origin. **F)** Dot plot showing normalized gene expression for canonical cell type markers in individual cell types detected within control PBMCs and unstimulated PBMCs treated with SOCE blockers.

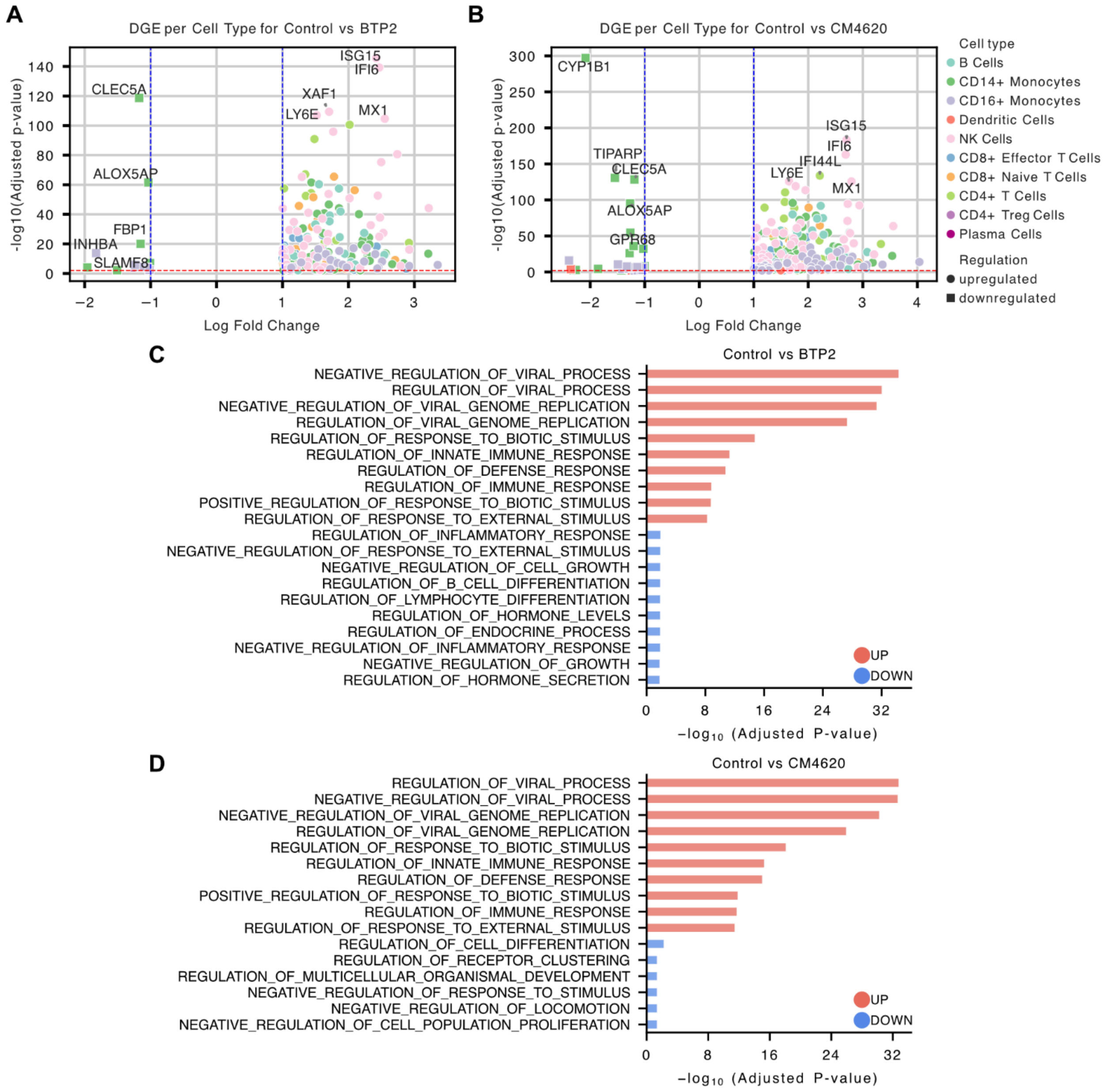


**Supplementary Figure 8: DGE of control PBMCs vs. PBMCs+BTP2 and PBMCs vs. PBMC+CM4620 per cell type and GO term analysis. A)** Volcano plot showing DGEA results for PBMCs (Control) vs. BTP2 for each cell type. **B)** Volcano plot showing DGEA results for PBMCs (Control) vs. CM4620 for each cell type. **A-B)** Colors correspond to the cell type and the shape to the direction of regulation. The blue line defines the +/-1.0 $\log_2$ fold-change threshold and the red line is set at 0.01 adjusted p-value. **C**) GO term analysis of all upregulated and downregulated genes of the cell type-split DGE analysis between Control and BTP2 treated unstimulated PBMCs. **D**) GO term analysis of all upregulated and downregulated genes of the cell type-split DGE analysis between Control and CM4620 treated unstimulated PBMCs. **C-D**) Only the DGE genes that met the +/-1.0 $\log_2$ fold-change threshold and had an adjusted p-value of 0.01 were considered.

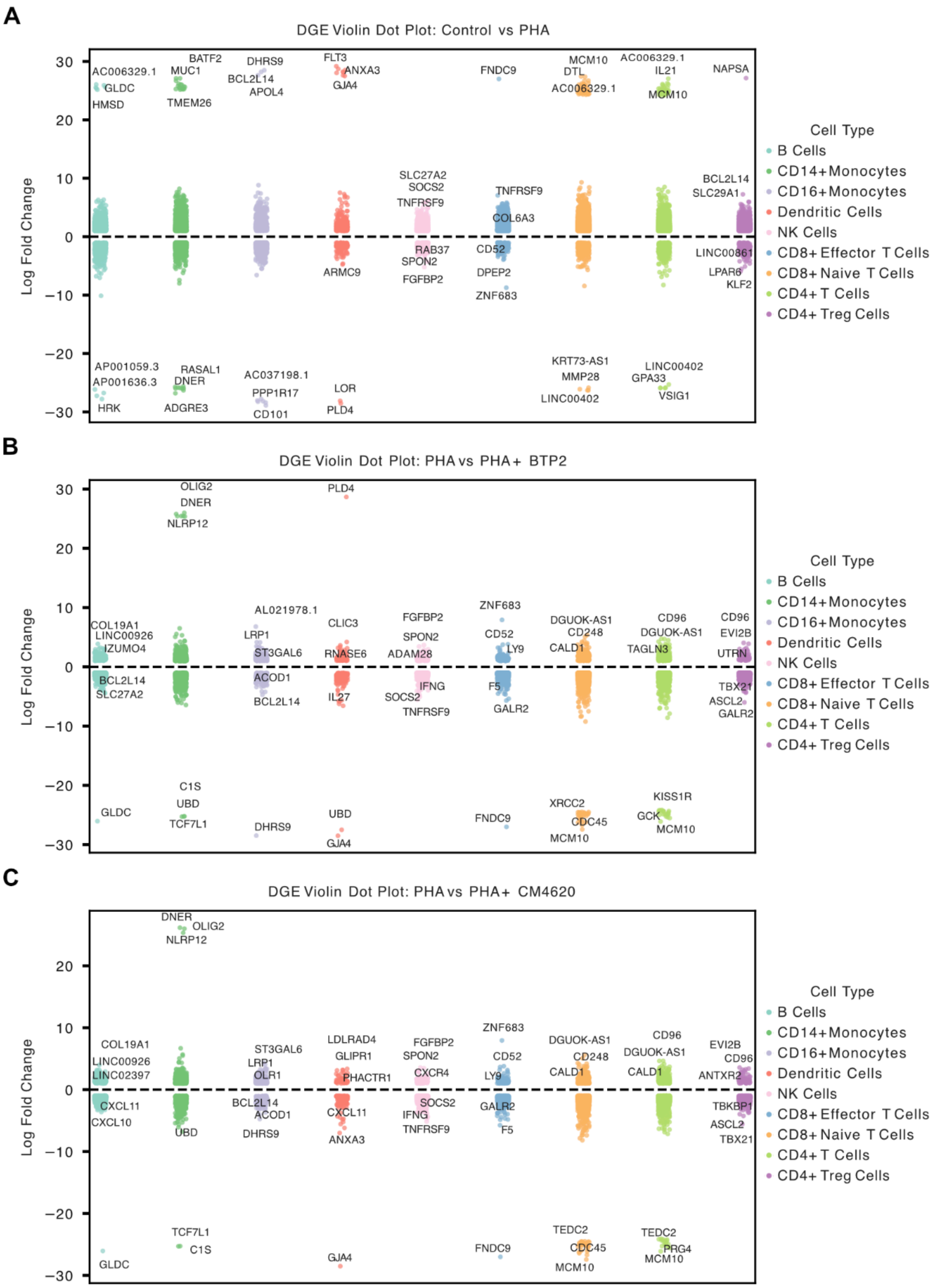


**Supplementary Figure 9: Differential gene expression analysis between treatment conditions for all cell types except plasma cells. A)** DGEA results for Control vs PHA. **B)** DGEA results for PHA vs PHA + BTP2. **C)** DGEA results for PHA vs PHA + CM4620. Only genes the top and bottom 3 genes that met the following thresholds were plotted: $\log_2$ fold-change > 1.0 or $\log_2$ fold-change < -1.0 and adjusted p-values < 0.01.

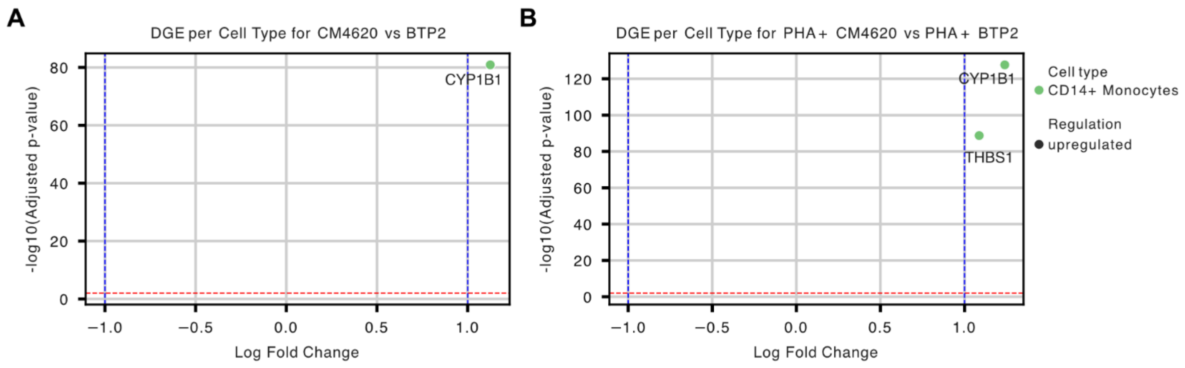


**Supplementary Figure 10: Differential gene expression analysis of PBMCs treated with SOCE blockers. A)** Volcano plot showing DGEA results for CM4620 vs. BTP2 for each cell type. **B)** Volcano plot showing DGEA results for PHA + CM4620 vs. PHA + BTP2 for each cell type. Colors correspond to the cell type and the shape to the direction of regulation. The blue line defines the +/-1.0 $\log_2$ fold-change threshold and the red line is set at 0.01 adjusted p-value.

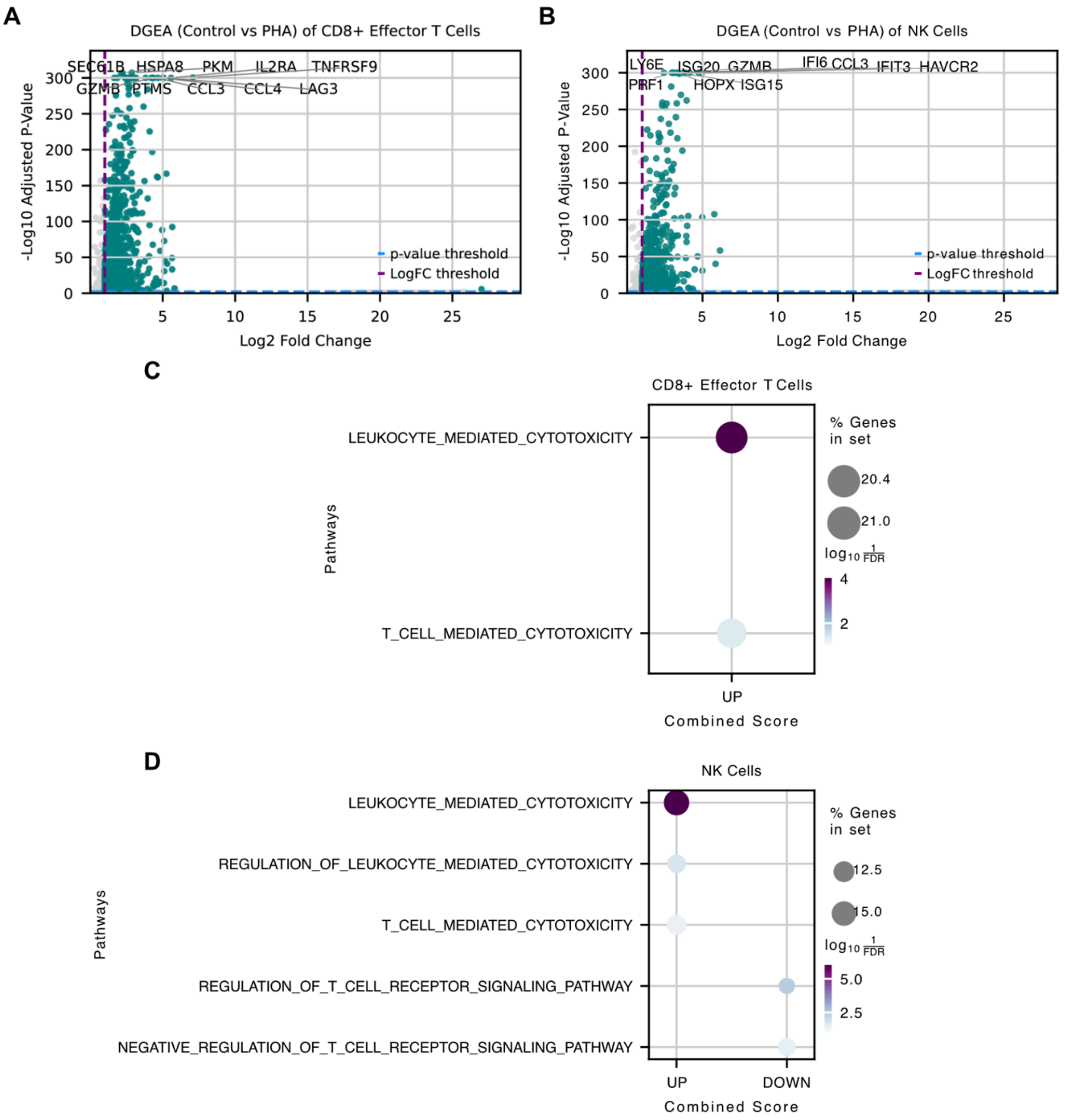


**Supplementary Figure 11: Differential gene expression and gene set enrichment analysis between conditions: Control PBMCs vs. PBMCs + PHA. A)** Volcano plot showing DGEA for CD8$^+$ Effector T cells in PHA-stimulated cells as compared to unstimulated controls. **B)** Volcano plot showing DGEA for NK cells in PHA-stimulated cells as compared to unstimulated controls. **A-B)** Pink dotted lines show the +1 log fold-change threshold, and the blue dotted lines show the adjusted p-value threshold of 0.01. Only the upregulated genes are plotted and only the top 10 upregulated gene names are shown. **C)** Dot plot showing top ontology terms enriched in DGEs for CD8$^+$ Effector T cells in PHA-stimulated PBMCs when compared to unstimulated controls. **D)** Dot plot showing top ontology terms enriched in DGEs for NK cells in PHA-stimulated PBMCs when compared to unstimulated controls. **C-D)** For gene set enrichment analysis only upregulated genes that met the following thresholds from the DGEA were used: $\log_2$ fold-change > 1.0 and adjusted p-values < 0.01.

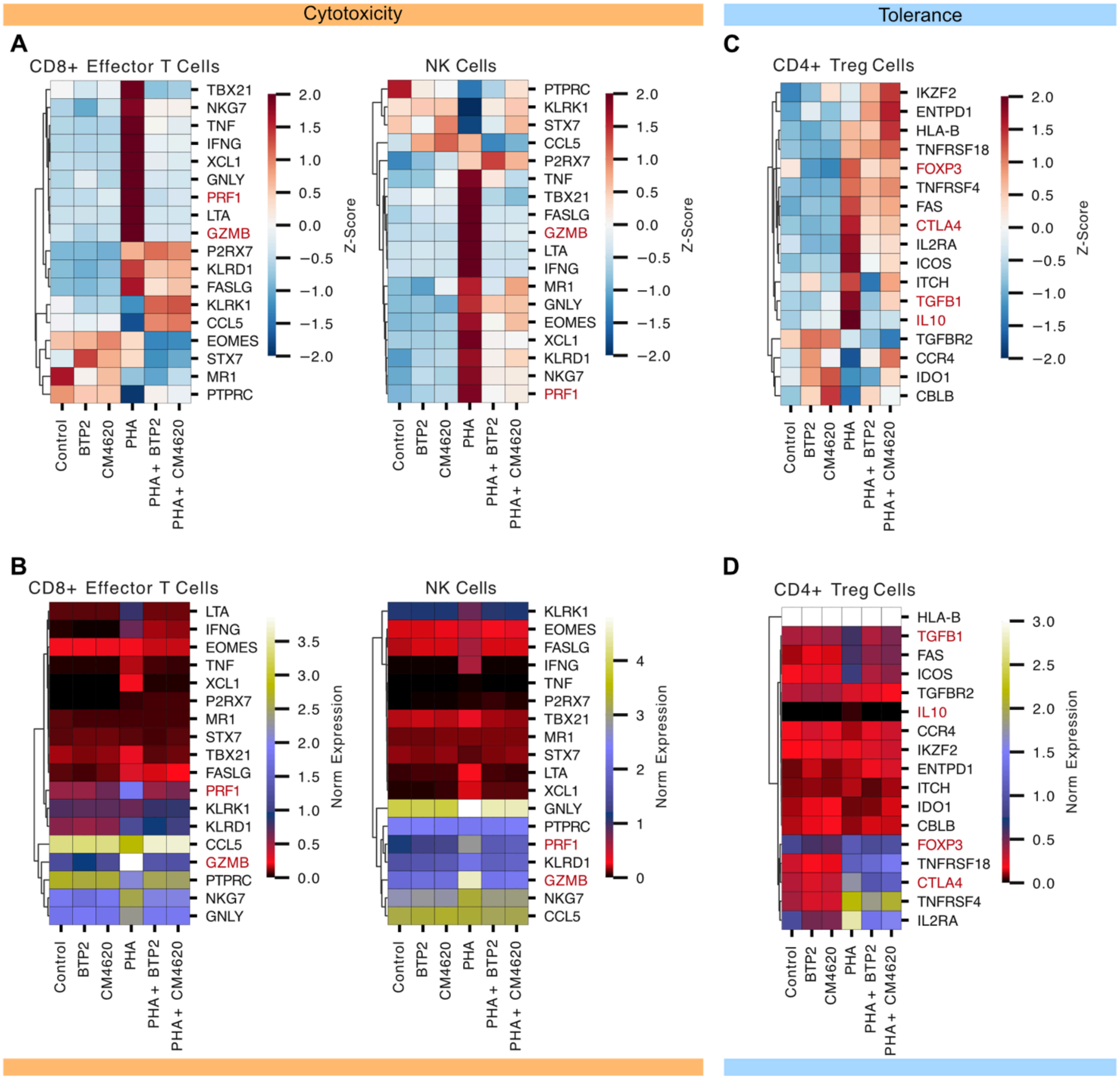


**Supplementary Figure 12**: **Gene expression of immune pathway related genes in T cells and NK cells. A)** Heatmaps showing z-scored normalized gene expressions of cytotoxic genes across different treatment conditions for CD8$^{+}$ Effector T cells and NK cells. **B)** Heatmaps showing the normalized gene expression for cytotoxic genes across different treatment conditions for CD8$^{+}$ Effector T cells and NK cells. **C)** Heatmaps showing z-scored normalized gene expressions of tolerance genes across different treatment conditions for CD4$^{+}$ T-reg cells. **D)** Heatmaps showing the normalized gene expressions for tolerance genes across different treatment conditions for CD4$^{+}$ T-reg cells. **A-D)** Red colored genes are the ones found in the RT-qPCR validation for cytotoxicity and tolerance.

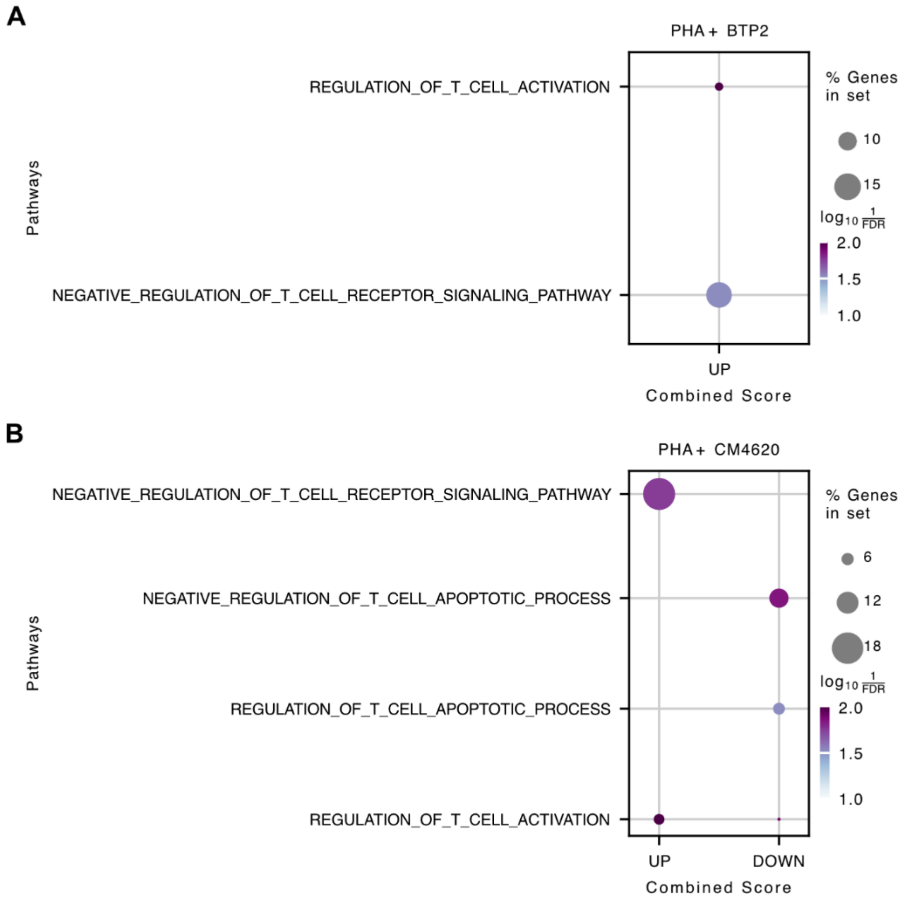


**Supplementary Figure 13**: **Gene set enrichment analysis for DGEA of CD4$^+$ T-reg cells between conditions A)** Dot plot showing ontology terms enriched in DGEA of Control vs. PHA + BTP2. **B)** Dot plot showing ontology terms enriched in Control vs. PHA + CM4620. For gene set enrichment analysis, only upregulated and downregulated genes that met the following thresholds from the DGEA were used: $\log_2$ fold-change > 1 for upregulated or $\log_2$ fold-change < -1 for downregulated and adjusted p-values < 0.01.
